# Ferries and environmental DNA: underway sampling from commercial vessels provides new opportunities for systematic genetic surveys of marine biodiversity

**DOI:** 10.1101/2021.05.05.442607

**Authors:** Elena Valsecchi, Antonella Arcangeli, Roberto Lombardi, Elizabeth Boyse, Ian M. Carr, Paolo Galli, Simon J. Goodman

## Abstract

Marine environmental DNA (eDNA) is an important tool for biodiversity research and monitoring but challenges remain in scaling surveys over large spatial areas, and increasing the frequency of sampling in remote locations at reasonable cost. Here we demonstrate the feasibility of sampling from commercial vessels (Mediterranean ferries) while underway, as a strategy to facilitate replicable, systematic marine eDNA surveys in locations that would normally be challenging and expensive for researchers to access. Sixteen eDNA samples were collected from 4 fixed sampling stations, and in response to 4 cetacean sightings, across three cruises undertaken along the 300 km ferry route between Livorno (Tuscany) and Golfo Aranci (Sardinia) in the Ligurian/Tyrrhenian Seas, June-July 2018. Using 12SrDNA and 16SrDNA metabarcoding markers, we recovered diverse marine vertebrate Molecular Operational Taxonomic Units (MOTUs) from teleost fish, elasmobranchs, and cetaceans. We detected sample heterogeneity consistent with previously known variation in species occurrences, including putative species spawning peaks associated with specific sea surface temperature ranges, and increased night time abundance of bathypelagic species known to undertake diel migrations through the water column. We suggest commercial vessel based marine eDNA sampling using the global shipping network has potential to facilitate broad-scale biodiversity monitoring in the world’s oceans.

## INTRODUCTION

Environmental DNA (eDNA) is an important tool to support biodiversity research and monitoring but challenges remain on how to scale eDNA surveys for assessments over large spatial areas, and to make it feasible to increase the frequency of sampling in remote locations at reasonable cost (Pawlowski et al., 2018). Such upscaling is important for generating high resolution biodiversity surveys needed for conservation planning, or impact assessments of human activities (Wetzel et al., 2018; Bani et al., 2020). Sampling design is often subject to logistical and financial constraints, which may mean data collection is opportunistic or has limited spatial and temporal resolution, introducing uncertainty into the interpretation of species presence/absence records (Menegotto and Rangel, 2018). However, understanding the environmental drivers of species distributions and interactions, requires systematic sampling over large spatial scales (Carstensen, 2014; Hale et al., 2018). eDNA has been suggested as a tool to improve the spatiotemporal resolution of biodiversity surveys and can offer the advantage of detecting communities of species from a single sample through the use of universal primer sets targeting taxa of interest (e.g. MiFish, Miya et al., 2015) and high throughput sequencing (HTS) (Stat et al., 2017). To date, studies in the marine environment have typically focused on detecting community differences at small spatial scales in coastal environments or comparing between regions using point-based samples (Port et al., 2016; Bakker et al., 2017; O’Donell et al., 2017; Yamamoto et al., 2017). Developing strategies to facilitate sampling in offshore pelagic environments could enhance the contribution of eDNA to large scale marine biodiversity surveys. This is particularly important for marine megafauna, where large distributions and dispersal capabilities of species mean that such taxa may have partial, low-resolution coverage from conventional techniques, while being a high priority for conservation planning due to their vulnerability and exposure to anthropogenic pressures (Hooker et al., 2011). In the case of marine mammals, eDNA studies have typically focused on single species assays (Foote et al., 2012; Baker et al., 2018; Hunter et al., 2018; Székely et al., 2021), or they have been found as serendipitous detections in metabarcoding surveys when using fish-specific primers (Closek et al., 2019). Developing metabarcoding approaches to reliably detect marine mammal eDNA in assays targeting marine vertebrates communities would increase efficiency and the scaling up of ecosystem level surveys and monitoring (Foote et al., 2012; Baker et al., 2018; Székely et al., 2021).

This study presents the first report, including novel sampling protocols, on using commercial marine ferry routes as a platform to systematically collect eDNA samples. Ferry routes allow sampling over the large spatial scales required for marine conservation planning and offer a cost-effective way for researchers to access remote offshore areas on a regular basis (Arcangeli et al., 2013; Matear et al., 2019). Since ferries typically follow set routes on routine schedules, they can be used for repeatable transect sampling over time (Arcangeli et al., 2013). Ferry-based sampling also allows the collection of samples at any time of the day (vessel schedule allowing), opening the opportunity to assess diel variation in marine community composition in environments that may not be accessible to smaller research vessels at night due to operating constraints. As a test case, we selected a route in the central Mediterranean basin between Livorno (Tuscany, Italy, Southern Ligurian Sea) and Golfo Aranci (Sardinia, Italy, Northern Tyrrhenian Sea) which crosses through the Pelagos Sanctuary for Mediterranean marine mammals (Notarbartolo□di□Sciara et al., 2008). We profile samples using two novel ‘universal’ marine vertebrate barcode markers employed here for the first time with marine environmental samples (Valsecchi et al., 2020), with the aim of investigating the potential of eDNA analysis to recover species community compositions known from conventional techniques, and variation in biodiversity patterns related to small scale spatial and temporal processes (such as diel variation and environmental factors such as bathymetry and sea surface temperature (Berry et al., 2019; Djurhuus et al., 2020). While we focus here on ferries as a test case, in principle the sampling approach can be deployed from any commercial vessel, opening up the possibility of sampling for marine eDNA across the global shipping network.

## MATERIALS AND METHODS

### Sample collection

Water samples were collected from the Mega Express III, a 212 m long vessel of the Corsica Ferries fleet, while underway along the route from Livorno (N 43° 33’ 36”, E 10° 18’ 0”) to Golfo Aranci (N 40° 59’ 24”, E 9° 37’ 12”) and on the return leg (hereafter LiGA route, see Figure 1). Samples were collected at four fixed sampling stations (FSS), selected to be evenly spaced along the route and to cover sites of ecological interest (e.g. where fishing effort is high, or in proximity to fish farms). Additional samples were taken opportunistically when cetaceans were sighted by a marine mammal observer team conducting visual surveys (see below). The latter, by necessity, were collected only during daylight hours, while fixed stations were sampled whenever the ferry crossed their location. Sampling was carried out during June and July 2018, on 3 return cruises, 15 days apart (18^th^-19^th^ June, 2^nd^-3^rd^ July, 16^th^-17^th^ July 2018; Supplementary Table S1). A sample naming convention was adopted as ‘LiGA*x*.*y*’ – where ‘*x*’ is the cruise number and ‘*y*’ – sampling station: 1, 2, 3, 4 for fixed stations; S1, S2 etc for sequential opportunistic sighting samples within cruises.

**Figure 1.**
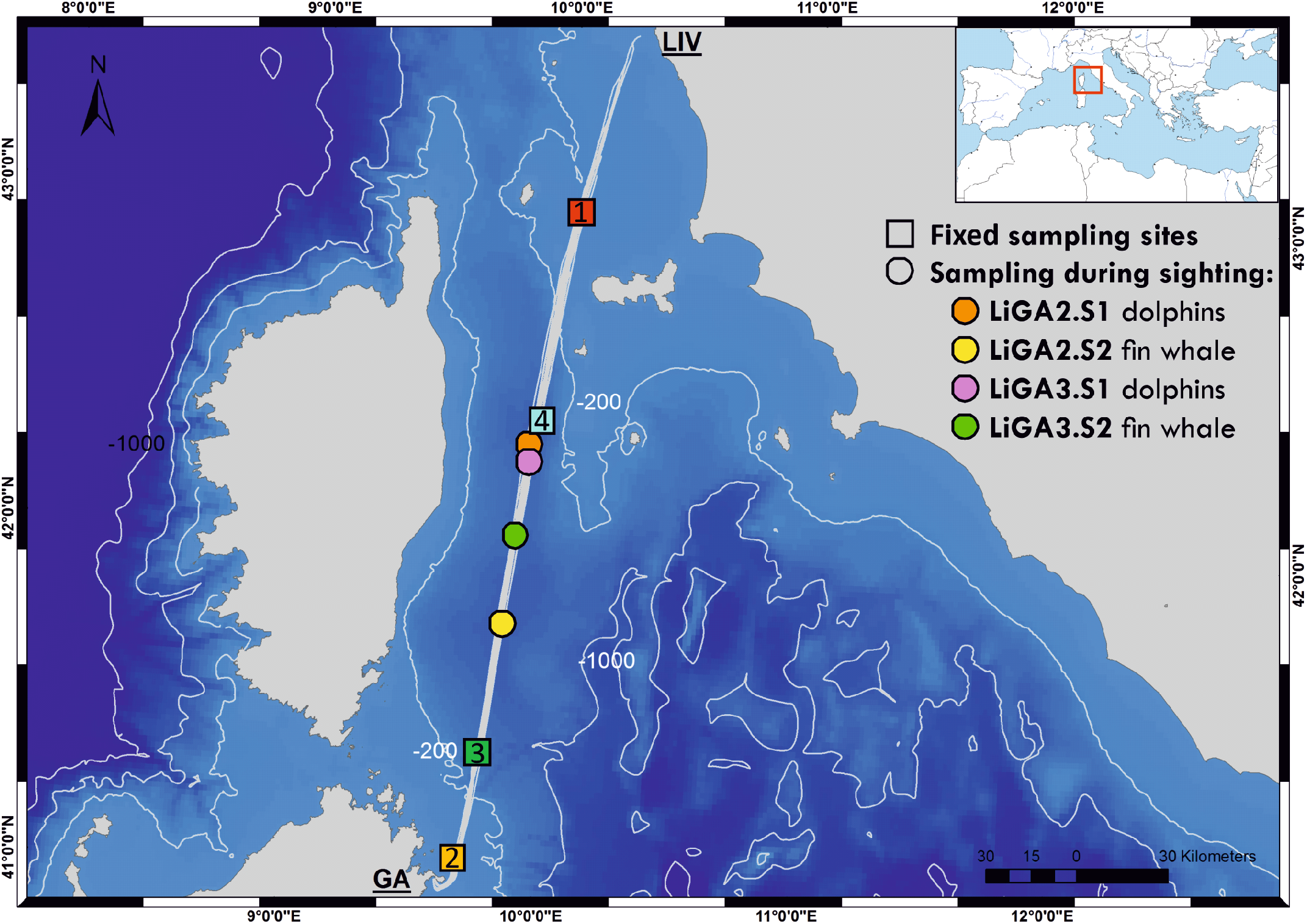
Bathymetric map of the northern Tyrrhenian Sea showing the ferry route between Livorno and Golfo Aranci, and the sampling sites for this study (coloured squares are fixed sites -LIGA1, 2, 3, 4- and circles correspond to opportunistic sampling associated with cetacean sightings). Inset map shows location of sampled route relative to the whole Mediterranean Sea. See Supplementary Table S1 for full details of sampling sites.

A total of 16 samples were collected: 12 from the FSSs and 4 during marine mammal sighting events (Figure 1 and Supplementary Table 1). Half (n=8) were collected during daylight and the remainder at night. Three fixed sampling stations (1, 3 and 4) were sampled with different light regimes over the 3 ferry trips according to the vessel schedule (FSS1: 2 day, 1 night; FSS3 and FSS4: 1 day, 2 night). All three FSS2 samples were collected at night.

### Cetacean observations by the FLT Mediterranean monitoring network

A marine mammal observer team from Istituto Superiore per la Protezione e la Ricerca Ambientale (ISPRA), part of the Fixed Line Transect (FLT) Mediterranean monitoring network (http://www.netccobams.com), travelled on each cruise. During daylight hours the ISPRA team recorded sightings of all cetacean species, and communicated encounters to the eDNA team in the vessel’s engine room, who immediately initiated water sample collection. The monitoring was conducted by professional observers positioned on both sides of the vessel bridge using a standard protocol based on distance sampling (ISPRA, 2015). A dedicated GPS was used to automatically record the survey track, marking the beginning/end points and the locations of sightings. Visual monitoring was only carried out when the wind strength was less than 3 on the Beaufort scale.

### Water sampling procedure

Water samples were collected from the ferry engine room, via a derivation pipe intercepting marine cooling water upstream of the engine. The water intake was located at 4.5 m below the sea surface. Before each sample was collected, the derivation pipe was opened allowing seawater to flow for 3-5 minutes to rinse the pipe with “local” water, thus removing residual water from the previous sample. A total of 13 litres of seawater was collected from each sampling site and was decanted directly from the derivation pipe into sterile foil laminated plastic “Bag-in-the-Box” (BiB) containers (supplied by KRCA, https://www.krca.it/; see Supplementary Figure S1). The BiB containers were subsequently used for sample storage and transportation.

The time to fill each BiB varied according to sea conditions and experience. On some occasions, the water coming out of the tap contained air which increased the time to fill the bag. Filling time ranged from 2 minutes to 10 minutes, with an average of 4 minutes 19 seconds (Supplementary Table S1). Considering that Mega Express III travels at a cruise speed of about 27.5 knots, samples were therefore collected while the ferry covered a mean distance of 2.27 nautical miles (3.66 km), with a minimum and maximum values of 1.05 nautical miles (1.7 km) and 5.27 nautical miles (8.49 km) respectively (Supplementary Table S1).

### Filtration

Most samples were filtered immediately after collection, unless multiple samplings for marine mammal sightings occurred in close succession, resulting in a slowdown of the filtration processing. To minimize this problem, we operated two portable vacuum pumps (Fisherbrand™ FB70155, Fisher Scientific) in parallel. In some instances, the water samples were filtered after returning to the laboratory in Milan. The time elapsed from sample collection to filtration ranged between 0 up to a maximum of 29 days, with a mean of 5 days.

Nitrocellulose filters with 0.8 μm, 0.45 μm and 0.22 μm pore sizes were used on aliquots of the same 13-litre sample to identify the best filtering protocol to use in future ferry eDNA sampling campaigns. For each membrane type, the volume of seawater passed through the filter was adjusted according to the onset of the filter’s pore saturation. Thus, 13 litre samples were subdivided into aliquots of 5, 5, and 3 litres filtered with 0.8 μm, 0.45 μm and 0.22 μm pore size nitrocellulose filters respectively, using a BioSart® 100 filtration system (Sartorius). Immediately after filtration, the membranes were folded on themselves (filtered-particle sides facing each other), wrapped in aluminium foil, labelled, and stored at - 20°C until DNA extraction.

Twelve of the sixteen samples were filtered in 3 aliquots with all 3 different membrane pore sizes. Four samples (LiGA1.4, LiGA3.1, LiGA3.4, LiGA3.S1), were filtered only with the 0.8 μm and the 0.45 μm membranes. Filtration duration varied among samples and filter type. The processing of 5 litres of marine water through the 0.8μm filtering membranes took from 1h 11’ to 8h 28’ (average 4h 52’); the same volume of water processed through the 0.45μm filters took from 1h 21’ to 10h 54’ (average 6h 39’), while the filtration of 3 litres of marine water through the 0.22μm took from 3h 40’ to 14h 14’ (average 8h 34’).

### DNA extraction, amplification and sequencing

DNA was extracted from filters between 0 and 31 days after filtration, with a mean of 14 days. Extractions were done using a DNeasy PowerSoil Kit^®^ (Qiagen), following the manufacturer’s protocol.

The target taxa for this study were all marine vertebrates, including cetaceans, so the DNA samples were amplified using MarVer1 (12S) and MarVer3 (16S) metabarcoding primers (Valsecchi et al., 2020). These primers were explicitly designed to increase efficiency for amplification of cetacean DNA whilst retaining the ability to amplify amplicons from teleosts, elasmobranchs, chelonians, and birds (Valsecchi et al., 2020). For each locus, seven independent PCR negative controls were included, which used sterile ultrapure water (Sigma Ltd) instead of an eDNA sample. The library for locus MarVer1 was sequenced in a 150bp paired-end lane, and locus MarVer3 in a 250bp paired-end lane, using an Illumina MiSeq sequencer at the University of Leeds Genomics Facility, St James’s Hospital, as described in (Valsecchi et al., 2020).

### Bioinformatic and data analyses

#### Initial quality filtering and annotation of amplicon sequences

The bioinformatics workflow was as described by Valsecchi *et al*. 2020 (Valsecchi et al., 2020). First, paired reads were screened for the presence of the expected primer and index sequence combinations to exclude off-target amplicons. Reads were then combined to generate insert sequences, and potential PCR duplicates and chimeric sequences were removed. The insert data was then aggregated to create a counts matrix containing the occurrence of each unique sequence in each sample. The taxonomic origin of each insert was determined by Blasting their sequence against a local instance of the GenBank NT database (Nucleotide, https://www.ncbi.nlm.nih.gov/nucleotide/). The level of homology of the insert to the hit sequence was noted, as was the species name of the hit sequence. The taxonomic hierarchy for each unique insert was generated by searching a local instance of the ITIS database (ITIS, https://www.itis.gov/) with the annotated GenBank species name. The count matrix and taxonomic hierarchy for all annotated unique sequences were then aggregated into values for equivalent Molecular Operational Taxonomic Units (MOTUs), by combining all inserts with a set homology (>=98%) to the GenBank hit at a specified taxonomic level (i.e., ‘Order’, ‘Family’, ‘Genus’ or ‘Species’), using bespoke software (available on request).

#### Relation between read counts and filter type/processing and sampling regimes

Pearson’s product moment correlation was used to evaluate read count recovery in relation to time elapsed from sampling to end of filtration, and time elapsed from filtration to extraction. Differences in read counts obtained between the 3 membrane pore-sizes, and for daylight vs night time sampling were assessed using Kruskal-Wallis tests. Subsequently, data from libraries with different filter sizes for the same sample were merged for further analyses.

#### Evaluation of potential contamination and ambiguous MOTU assignments

Following an approach similar to that suggested by (McKnight et al., 2019), MOTUs attributable to potential contamination were identified by comparing the ratio of mean read counts for each MOTU in marine samples, versus the mean read counts of the same MOTU in the 7 PCR negative controls. MOTUs with a ratio of less than 20:1 (Sample:Control) were excluded, except for 10 cases where the ratio was greater than 5:1, and the MOTU was not related to any species previously handled in the analysing laboratory, and/or the ratio for the alternate marker was greater than 20. Excluded MOTUs were primarily amplicons attributable to taxa present in environmental samples from Genoa aquarium which were analysed in the same sequencing runs (Valsecchi et al., 2020). In addition, MOTUs from humans, cow, pig, dog, chicken, turkey, and other taxa which were related to control DNAs from (Valsecchi et al., 2020) were removed. For MarVer1 and MarVer3, 8.6 % and 3.6 % of reads respectively were excluded under these criteria (see Table 1; Supplementary Tables S3, S4).

**Table 1.** Number of reads and MOTUs for across main taxonomic division for MarVer1 and MarVer3 markers. Excluded refers to reads and MOTUs excluded as potential contamination (see Supplementary Tables S3 and S4 for taxon-sample read counts).

|  | MarVer1 |  |  | MarVer3 |  |  |
| --- | --- | --- | --- | --- | --- | --- |
|  | Percentage total |  |  | Percentage total |  |  |
|  | Reads | reads | MOTUs | Reads | reads | MOTUs |
| **Total annotated reads** | 606544 | - | 93 | 1021207 | - | 129 |
| **Excluded** | 52106 | 8.591 | 40 | 36777 | 3.601 | 44 |
| **Teleost** | 553801 | 91.304 | 45 | 979340 | 95.900 | 76 |
| **Cetacean** | 383 | 0.063 | 4 | 4111 | 0.403 | 4 |
| **Elasmobranch** | 240 | 0.040 | 2 | 340 | 0.033 | 1 |
| **Aves** | 14 | 0.002 | 2 | 0 | 0.000 | 0 |
| **Invertebrate** | 0 | 0.000 | 0 | 639 | 0.063 | 4 |

Amplicon sequences for MOTUs annotated as species not recorded for the Mediterranean (based on the Fishbase catalogue (www.fishbase.se)) were evaluated individually against related sequences in Genbank according to the scheme shown in Supplementary Figure S2. Such MOTUs were then either reclassified as derived from a Mediterranean taxon, or highlighted as a potential newly described record or introduced non-indigenous species (NIS; see Supplementary Table S2).

#### Analyses of community composition and sample diversity in relation to ecological and environmental variables

Summaries and visualisations of read counts for different taxonomic levels were generated using the R package ‘Phyloseq’ (McMurdie and Holmes, 2013). Supporting data for environmental variables, such as sea surface temperature (SST), chlorophyll concentration (CPH), salinity and bathymetry, covering sampling locations and dates (Supplementary Table S1) were obtained from the EU Copernicus Marine Service. Habitat type and trophic levels for fish species were downloaded from www.fishbase.se.

Associations between read counts of specific MOTUs, or other summary statistics such as measures of Alpha diversity for samples, and environmental variables were assessed through Pearson’s product-moment correlation.

## RESULTS

### Read counts in relation to sample categories

After quality filtering, a total of 606,544 and 1,021,207 reads for MarVer1 and MarVer3 respectively, were assigned to taxa with Genbank references (Table 1). No significant correlation was found between the number of reads recovered from each sample and the time span from sampling to filtration for MarVer1 (*t* = -0.0622, df = 42, *p* = 0.951, cor = -0.0096), but a weak negative correlation was detected for MarVer3 (*t* = -2.3787, df = 40, *p* = 0.0222, cor = -0.3520). Most samples (95.5%) were filtered within 8 days from collection and, within that time window, sample read counts were evenly distributed (Supplementary Figure S3a). A negative correlation was observed for read counts and time from filtration to extraction for MarVer1 (*t* = -2.8145, df = 41, *p* = 0.0075, cor = -0.4024), and a positive association for MarVer3 (*t* = 2.5137, df = 39, *p* = 0.0162, cor = 0.3734). However, read counts were relatively evenly distributed overall, and the significant correlations may be driven by outlying values (Supplementary Figure S3b). No difference in mean read counts among filter pore sizes was detected for either locus (Supplementary Figure S3c; MarVer1, Kruskal-Wallis χ^2^ = 0.2019, df = 2, *p* = 0.9039; MarVer3, Kruskal-Wallis χ^2^ = 2.2511, df = 2, *p* = 0.3245). A significantly higher mean number of reads were recorded in night-time versus day-light samples (Supplementary Figure S3d) for both MarVer1 (Kruskal-Wallis χ^2^ = 12.34, df = 1, *p* = 0.0004) and MarVer3 (Kruskal-Wallis χ^2^ = 4.6684, df = 1, *p* = 0.0307). Finally, no significant correlation was detected between read count and sample collection track length (km), MarVer1: t = -0.4867, df = 14, *p* = 0.634, cor = -0.1290; MarVer3: t = 1.8967, df = 14, *p* = 0.07869, cor = 0.4521; and there was no significant difference in mean read count per sample among cruises (Supplementary Figure S4; MarVer1: Kruskal-Wallis χ^2^ = 1.7831, df = 2, *p* = 0.41; MarVer3: Kruskal-Wallis χ^2^ = 1.4706, df = 2, *p* = 0.4794).

### Summary of taxonomic assignments

More than 90 % of reads were attributed to bony fish species, with the second most represented group being cetaceans, followed by elasmobranchs and birds (Table 1, Figure 2). Combined across both loci, a total of 92 unique teleost MOTUs from 31 families were detected, comprising 45 by MarVer1 and 77 by MarVer3, with 30 detected by both markers. Other vertebrate taxa detected included 4 cetacean MOTUs (see below), 2 bird MOTUs (MarVer1 only: Scopoli’s shearwater (*Calonectris diomedea*) and herring gull (*Larus argentatus*)), and 2 ray MOTUs (devil fish, *Mobula mobular* and pelagic stingray, *Pteroplatytrygon violacea*; MarVer1 only). MarVer3 also recovered a small number of amplicons from 4 invertebrate MOTUs (3 hydrozoans - *Aglaura hemistoma, Geryonia proboscidalis, Liriope tetraphylla*, and a *Phascolosoma* peanut worm species), representing 0.06% of total reads.

**Figure 2.**
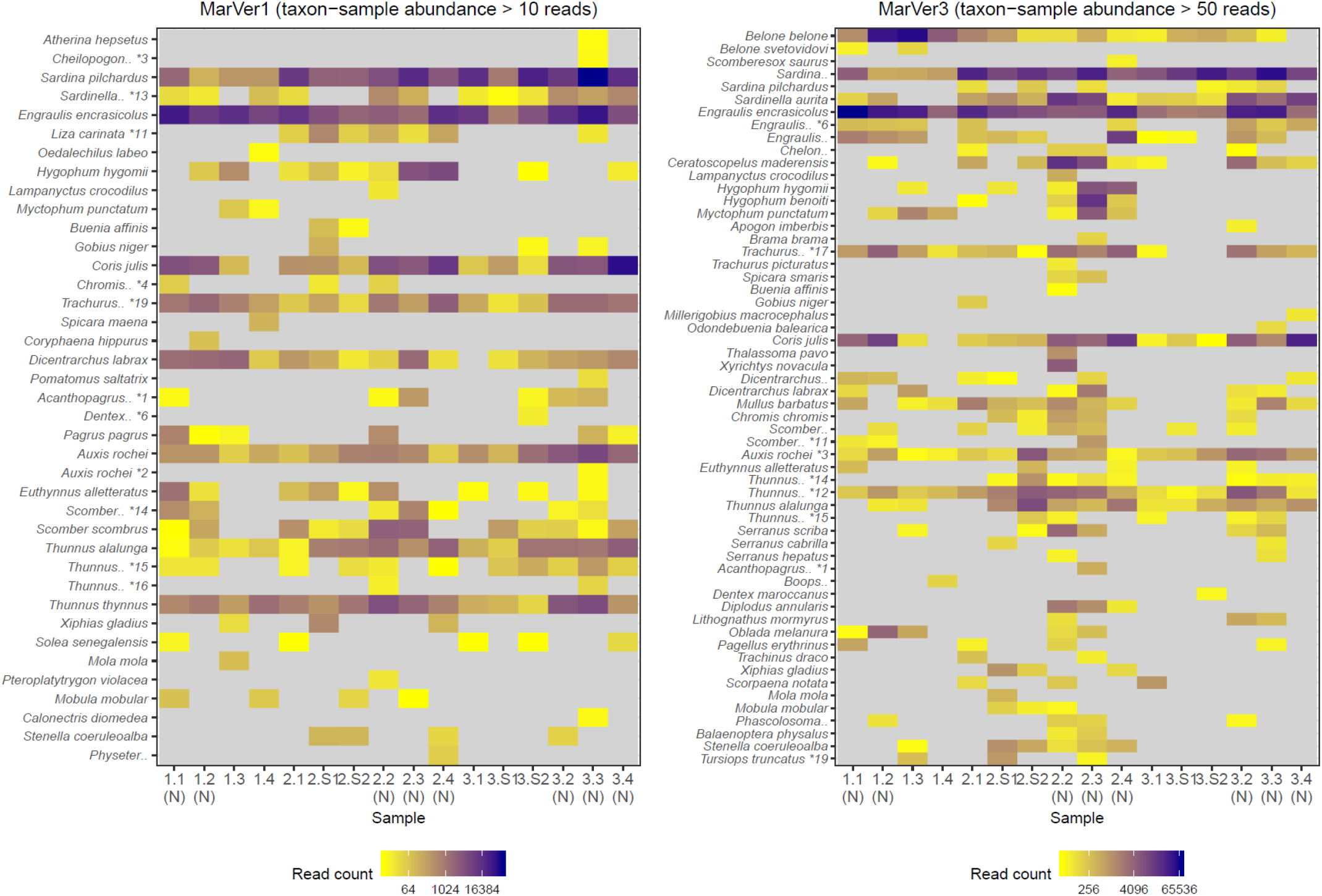
A heatmap of MOTU read counts in each sample recovered for the MarVer1 and MarVer3 loci. MOTU abundance is indicated by the colour scale. Rare MOTU occurrences with taxon-sample abundances less than 10 and 50 reads respectively were excluded from the figure to improve readability. (N) – night-time sample, (S) – cetacean sighting sample. MOTUs annotations with ‘*’ were adjusted as described in Supplementary Table S2. Full MOTU counts are given in Supplementary Tables S3 and S4.

Initially, 18 MOTUs for both loci were attributed to non-Mediterranean species, but most were subsequently resolved to Mediterranean taxa (Supplementary Table S2). These represent cases either where there is no reference sequence available for the corresponding species, or where variation among haplotypes is less than the 2% divergence threshold used to amalgamate MOTUs. These included several tuna MOTUs (particularly *Thunnus albacares* which was detected at high abundance by both markers), whereas only 2 tuna species (*Thunnus alalunga, Thunnus thynnus*) are considered Mediterranean residents. A MOTU assigned to the non-resident *Trachurus japonicus*, was also detected at high abundance by both markers. The Atlantic (*Trachurus trachurus*) and Mediterranean (*Trachurus mediterraneus*) horse mackerels are normally found in the Mediterranean, but for the latter there is no reference sequence deposited, and the Atlantic species *Trachurus trachurus* differs only by 1 and 4 bp from *Trachurus japonicus* sequence for MarVer1 and MarVer3 respectively. Similar cases occurred for cetaceans - amplicons annotated as the non-resident *Tursiops aduncus* were detected by MarVer3, which were taken as originating from *T. truncatus* (from which it differs by 1 variable site), and reads for *Stenella frontalis* and *S. longirostris* were recovered by MarVer3 (2 variable sites) and MarVer1 (1 variable site) respectively, but we assume are derived from *S. coeruleoalba*. For other MOTUs, where a non-resident could be unambiguously associated with a single resident congeneric, we consider them as originating from that resident species, otherwise, we consider them unresolved at the genus level. Ultimately 4 and 3 MOTUs remained unrelated to known Mediterranean taxa for MarVer1 and MarVer3 respectively. However, these instances were all low abundance amplicons accounting for only 110 (0.0181 %) and 38 (0.0037 %) reads for MarVer1 and MarVer3 respectively. None of these MOTUs scored more than 10 reads in a single sample. For MarVer1 the species were: *Cololabis saira* (6 reads), *Dentex tumifrons* (56 reads), *Engraulis japonicus* (47 reads) and *Gasterochisma melampus* (1 read); and for MarVer3: *Cyclothone atraria* (31 reads), *Dentex canariensis* (6 reads) and *Lutjanus fulvus* (1 read). Given the low abundance of these MOTUs, further interpretation would be speculative as to whether they represent artefacts, possible detections of new alien species or novel resident taxa. However, if future studies corroborate the incidence of these taxa, then our data should be reviewed as early detection of the species.

### Fish community composition

The two markers showed high consistency in the overall relative abundances of detected teleost taxa, with 8 genera in common among the 10 most abundant for each marker (Table 2). Sample profiles were dominated by anchovy (*Engraulis* spp.) and sardine (*Sardina* spp.) MOTU reads, accounting for 32.8% and 33.8%, and 32.2% and 24.1% of teleost reads for MarVer1 and MarVer3 respectively. Overall the 10 most abundant genera accounted for 98.8% and 94.5% of reads for MarVer1 and MarVer3. Intermediate trophic level predatory fish (trophic level 3) were the most commonly detected species type (greater than 75% total reads; Figures 2, 3, 4), but top-level predators (trophic level 4 in Figure 3) including Garfish (*Belone* spp.; detected by MarVer3 only), Tuna (*Thunnus* spp), and Bullet/Frigate tuna (*Auxis* spp.) were also among the top 10 most abundant taxa. Amongst the less abundant MOTUs, detections of eDNA from other large predatory fish included swordfish (*Xiphias gladius*), and ocean sunfish (*Mola mola*), which were found by both markers in multiple samples and from different cruises. With respect to habitat type (as defined in www.fishbase.se), MOTUs from pelagic-neritic species were most abundant (greater than 60 % reads), followed by pelagic-oceanic, reef associated and bathypelagic (Figure 3).

**Table 2.**
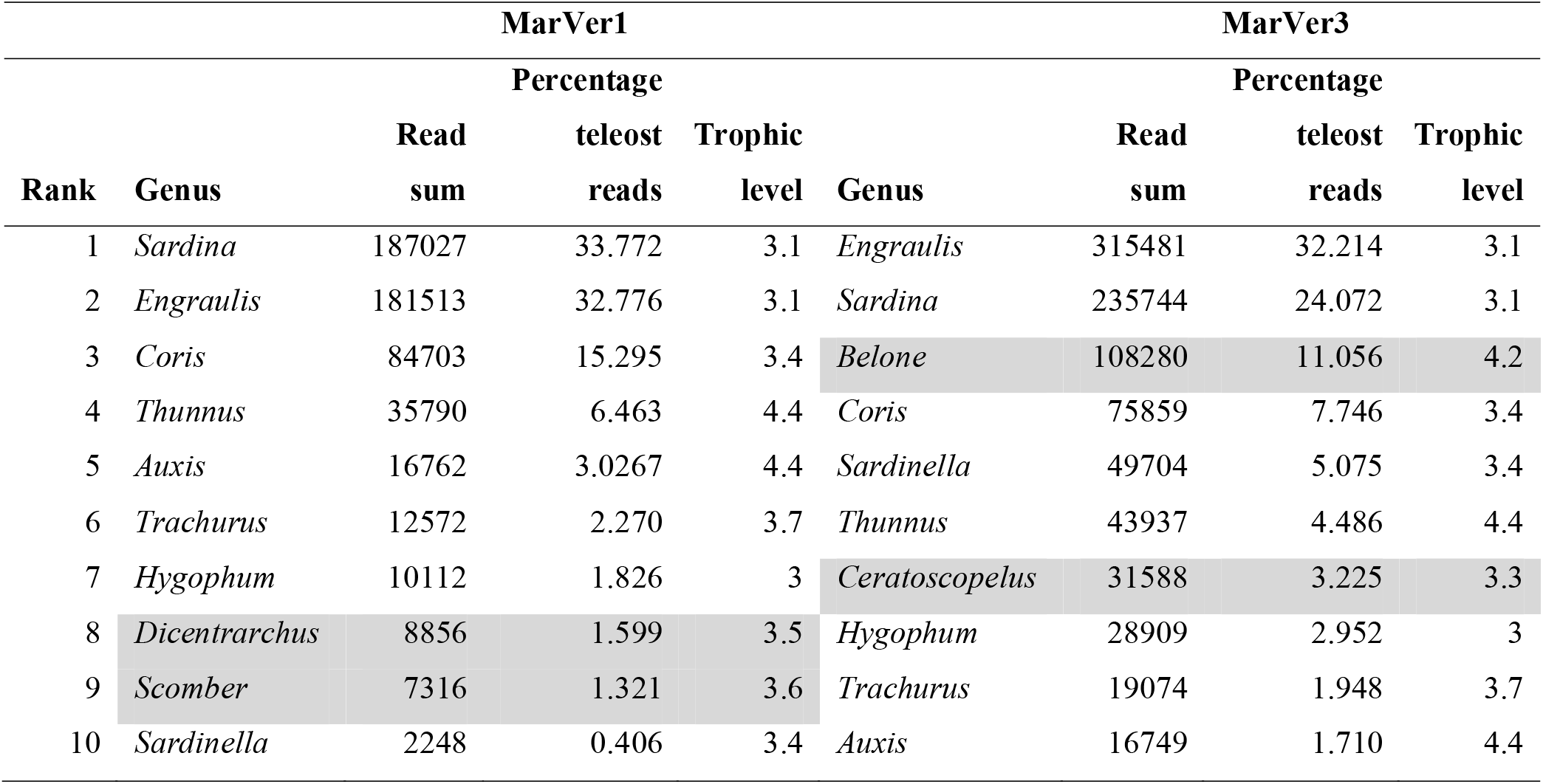
Top 10 most abundant combined MOTU genera by read count for MarVer1 and MarVer3. Shaded taxa indicate genera not shared in top 10 by both markers. Trophic level taken from catalogue of Mediterranean fish at www.fishbase.se, accessed December 2020.

**Figure 3.**
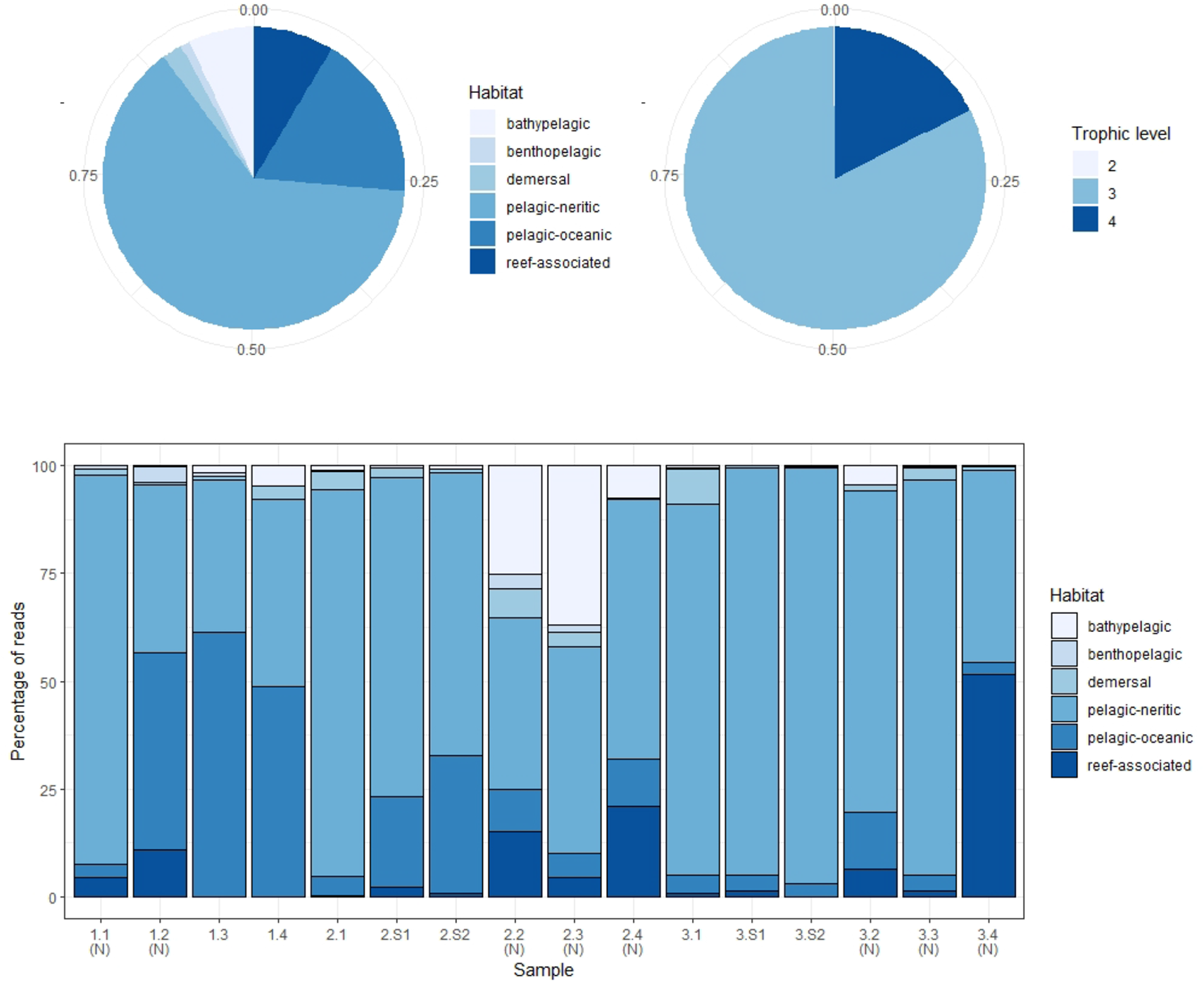
Summary of proportions of species habitat types and trophic levels for MarVer3 MOTUs, for the overall sample (pie charts), and individual samples (barchart). (N) – night-time sample, (S) – cetacean sighting sample. Species habitat type and trophic level definitions taken from www.fishbase.se.

**Figure 4.**
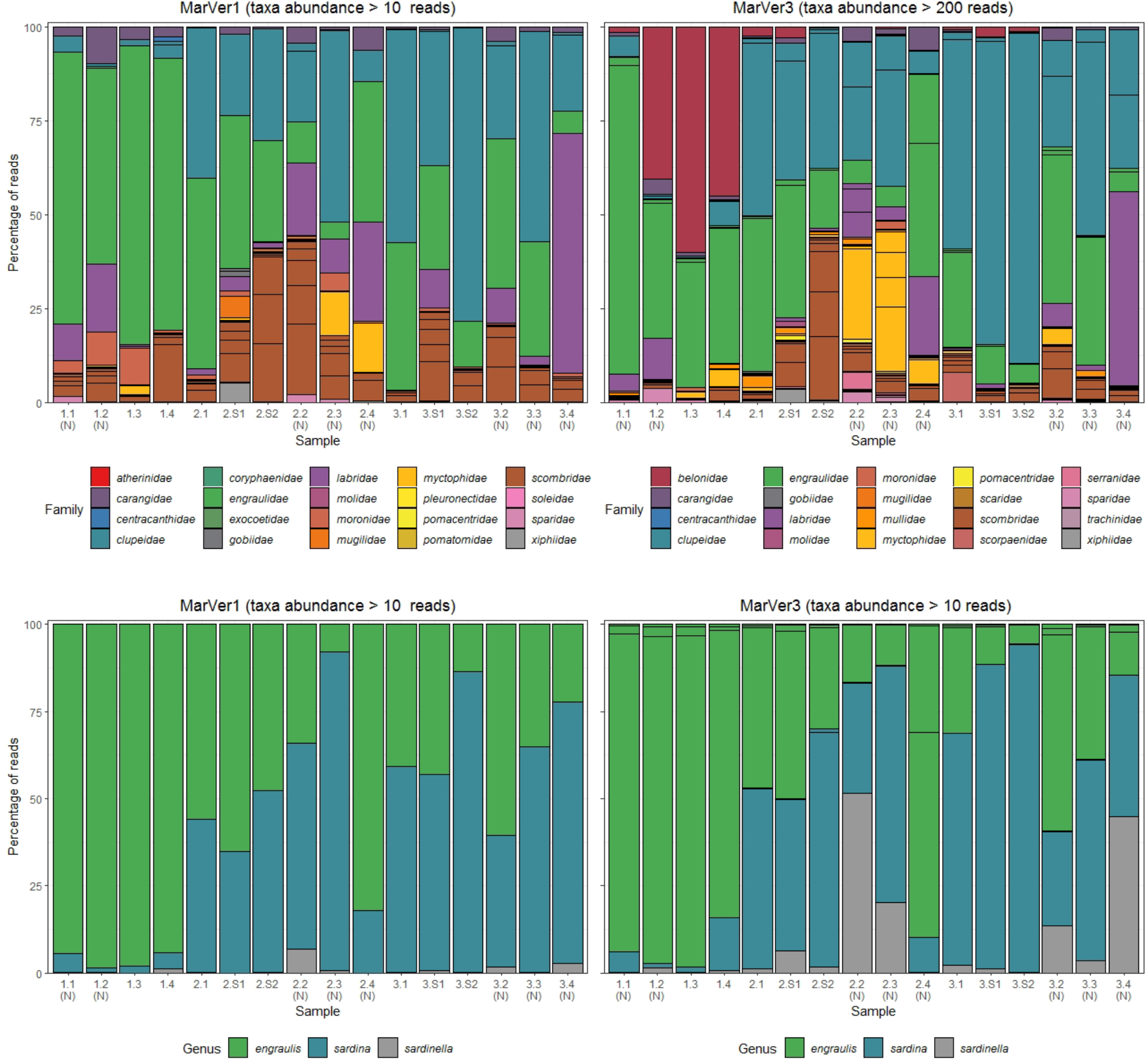
Barcharts showing proportions of the 20 most abundant teleost fish families (upper), and anchovy and sardine/sardinella genus MOTUs in each sample for MarVer1 and MarVer3. (N) – night-time sample, (S) – cetacean sighting sample.

Species compositions showed considerable spatiotemporal variation across samples. For example, *Belone* MOTUs were detected at high abundance in fixed station samples LiGA1.2-1.4 by MarVer3, but were rare at the same locations for cruises 2 and 3, sampled 15 and 30 days later respectively (Figures 2, 4). Mediterranean rainbow wrasse (*Coris julis*; Family Labridae) was detected at high abundance by both markers at fixed station 4 for cruise 2 and 3 (samples 2.4 and 3.4), but had lower abundance in other samples. However, the relative patterns of abundance for the most common taxa were consistent between the two loci (Figures 2, 4).

The relative abundances of anchovy (*Engraulis* spp.) and sardine/sardinella (*Sardina/Sardinella* spp.) MOTUs showed a reciprocal occurrence pattern across different samples for both markers, such that high abundances of *Engraulis* MOTUs corresponded with low abundances of sardine/sardinella and vice versa (Figures 2, 4). *Sardina* showed low abundance (less than 10%) in samples for Cruise 1 (mid-June), but predominated in 8 of 12 samples from Cruises 2 and 3 (July). These patterns showed a significant association with sea surface temperatures across a range of 22.7°C to 26°C, with *Engraulis* showing a negative relationship, and *Sardina* a positive correlation (Table 3, Supplementary Figure S5).

**Table 3.**
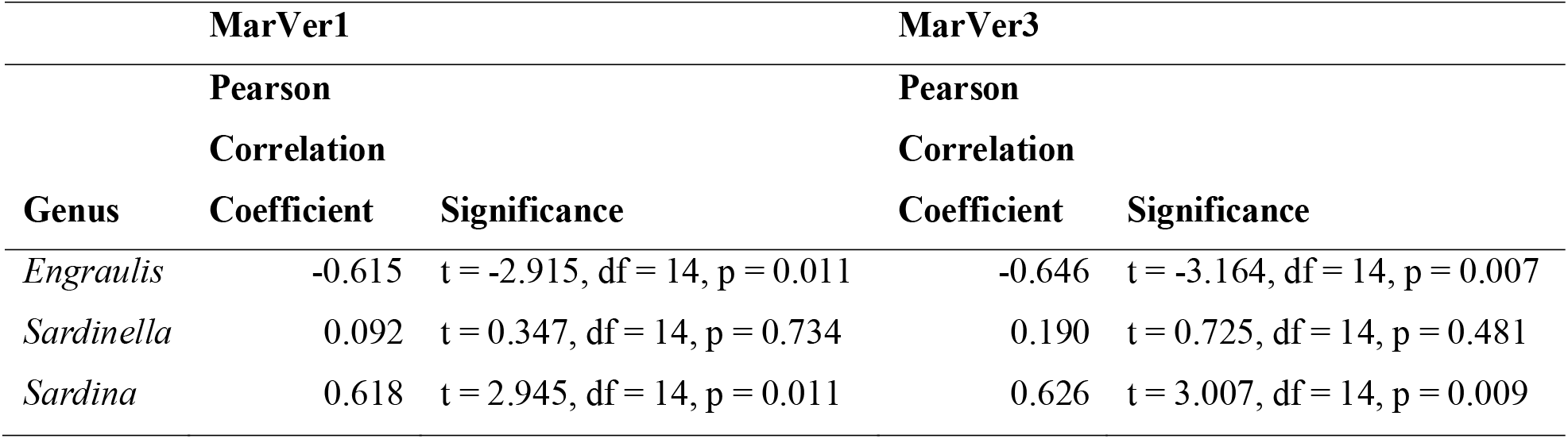
Pearson product moment correlation statistics for comparisons of relative Clupeidae genera proportions in samples versus estimated sea surface temperature at time of sampling, for each locus (See Supplementary Figure S5).

Both markers also detected eDNA from several lantern fish species (Myctophidae family) including *Ceratoscopelus maderensis, Lampanyctus crocodilus, Hygophum hygomii, Hygophum benoiti*, and *Myctophum punctatum*. These are bathypelagic species which undertake a vertical diel migration into the epipelagic zone at night. Read abundances for these and other bathypelagic species showed increases in several of the nocturnal sample collections, notably in cruise 2, which coincided with a near full moon (88% full, Figure 2 and Table S1).

### Detection of cetofauna

Traces of cetacean eDNA were detected for 8 of the 16 sample sites with MarVer1 and from all sites (fixed and sightings) with MarVer3 (Figure 5, Supplementary Tables S3 and S4). The eDNA signals were weak, with 97 % and 87 % of species-sample combinations for MarVer1 and MarVer3 respectively yielding fewer than 100 reads. Both primer sets detected amplicons attributable to bottlenose dolphins (*Tursiops truncatus*), striped dolphins (*Stenella coeruleoalba*), and the fin whale (*Balaenoptera physalus*). MarVer1 also detected sperm whale (*Physeter macrocephalus*) eDNA in the nocturnal sample LiGA2.4 (03/07/2018, h03:09). All four species had at least one detection with a minimum of 50 read counts, for one or both markers. Reads for striped dolphin were the most abundant for both markers, with a maximum per sample abundance of 120 for MarVer1 and 1098 for MarVer3.

**Figure 5.**
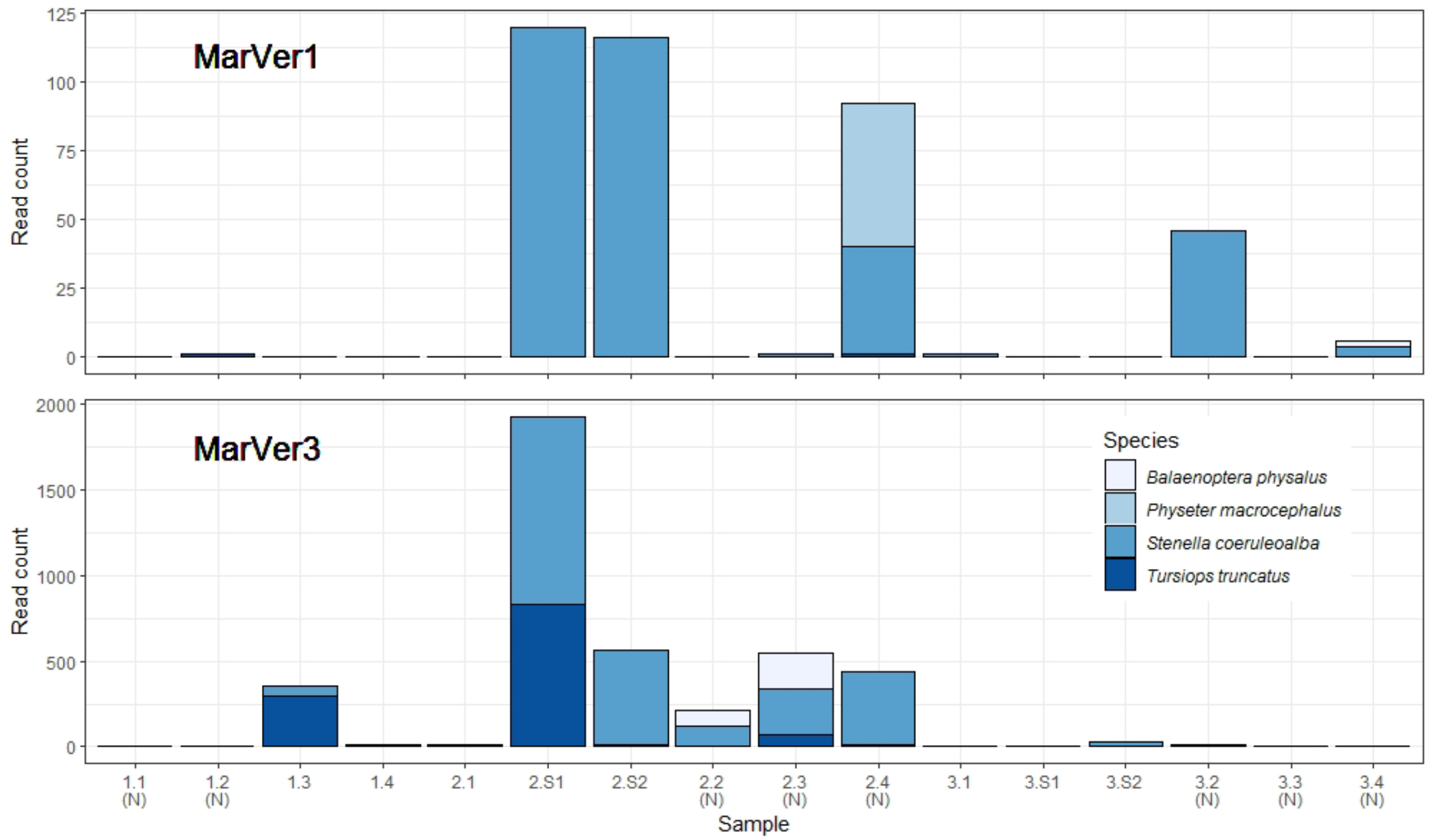
Barcharts showing read counts for cetacean MOTUs in each sample for MarVer1 and MarVer3. (N) – night-time sample, (S) – cetacean sighting sample.

Cetacean eDNA was detected in all four visual sighting samples for MarVer3 and at LiGA2.S1 and LiGA2.S2 for MarVer1. In these two samples, both markers recovered more than 100 reads and matched the species detected by visual observations. Mean read counts were higher for visual sighting samples than fixed stations for both loci, and for day-time versus night-time when including visual sighting samples. However, when excluding sighting samples, mean night-time abundances were higher.

### Intra and inter-sample diversity

Alpha diversity measures for MOTUs (Shannon and Inverse Simpson; Table 4) were highest for both markers in samples from Cruise 2, and fixed station 2 had the highest diversity on each Cruise. Mean diversities for night time samples were higher than those collected in daylight. Mean diversity measures varied significantly across cruises for the Inverse Simpson measure (MarVer1: Kruskal-Wallis χ^2^ = 7.3419, df = 2, p = 0.02545; MarVer3: Kruskal-Wallis χ^2^ = 7.989, df = 2, p = 0.01842), but differences in means for other comparisons were not significant, nor were correlations between sample diversities and read count, sea surface temperature, chlorophyll concentration, and collection track length.

**Table 4.**
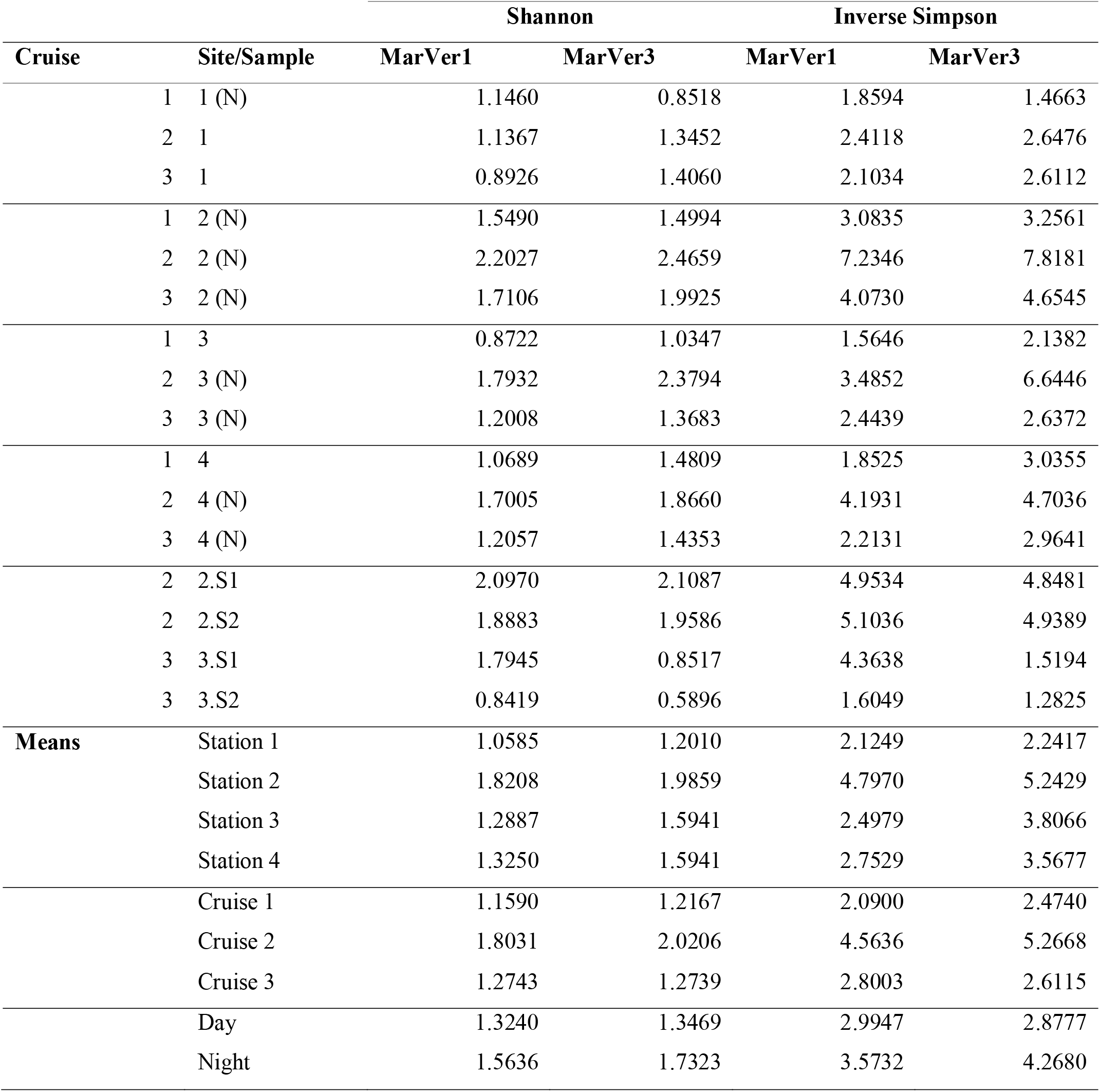
Sample alpha diversity measures (Shannon, Inverse Simpson) and means across fixed stations (1, 2, 3, 4), Cruises, and day/night. (N) – night sample; S1, S2 – cetacean sighting samples.

| Cruise | Site/Sample | Shannon |  | Inverse Simpson |  |
| --- | --- | --- | --- | --- | --- |
|  |  | MarVer1 | MarVer3 | MarVer1 | MarVer3 |
| 1 | 1 (N) | 1.1460 | 0.8518 | 1.8594 | 1.4663 |
| 2 | 1 | 1.1367 | 1.3452 | 2.4118 | 2.6476 |
| 3 | 1 | 0.8926 | 1.4060 | 2.1034 | 2.6112 |
| 1 | 2 (N) | 1.5490 | 1.4994 | 3.0835 | 3.2561 |
| 2 | 2 (N) | 2.2027 | 2.4659 | 7.2346 | 7.8181 |
| 3 | 2 (N) | 1.7106 | 1.9925 | 4.0730 | 4.6545 |
| 1 | 3 | 0.8722 | 1.0347 | 1.5646 | 2.1382 |
| 2 | 3 (N) | 1.7932 | 2.3794 | 3.4852 | 6.6446 |
| 3 | 3 (N) | 1.2008 | 1.3683 | 2.4439 | 2.6372 |
| 1 | 4 | 1.0689 | 1.4809 | 1.8525 | 3.0355 |
| 2 | 4 (N) | 1.7005 | 1.8660 | 4.1931 | 4.7036 |
| 3 | 4 (N) | 1.2057 | 1.4353 | 2.2131 | 2.9641 |
| 2 | 2.S1 | 2.0970 | 2.1087 | 4.9534 | 4.8481 |
| 2 | 2.S2 | 1.8883 | 1.9586 | 5.1036 | 4.9389 |
| 3 | 3.S1 | 1.7945 | 0.8517 | 4.3638 | 1.5194 |
| 3 | 3.S2 | 0.8419 | 0.5896 | 1.6049 | 1.2825 |
| **Means** | Station 1 | 1.0585 | 1.2010 | 2.1249 | 2.2417 |
|  | Station 2 | 1.8208 | 1.9859 | 4.7970 | 5.2429 |
|  | Station 3 | 1.2887 | 1.5941 | 2.4979 | 3.8066 |
|  | Station 4 | 1.3250 | 1.5941 | 2.7529 | 3.5677 |
|  | Cruise 1 | 1.1590 | 1.2167 | 2.0900 | 2.4740 |
|  | Cruise 2 | 1.8031 | 2.0206 | 4.5636 | 5.2668 |
|  | Cruise 3 | 1.2743 | 1.2739 | 2.8003 | 2.6115 |
|  | Day | 1.3240 | 1.3469 | 2.9947 | 2.8777 |
|  | Night | 1.5636 | 1.7323 | 3.5732 | 4.2680 |

A hierarchical cluster analysis of Bray-Curtis distances among samples (Supplementary Figure S6) identified all the sighting samples clustering together for MarVer3, and other paired clusters of night-time, or within-cruise samples were observed. However, the study was not designed to test for associations with environmental or ecological variables, so such patterns should be treated cautiously. Nonetheless, this provides some evidence for heterogeneity and structure in species composition among samples.

## DISCUSSION

In this study we evaluated the feasibility of using large commercial vessels, such as ocean-going ferries, as a platform for eDNA sampling to support marine biodiversity studies. Metabarcoding results from mitochondrial markers targeting vertebrates recovered diverse community profiles matching known Mediterranean biodiversity, including teleost fish, elasmobranchs, birds and cetaceans. We also detected inter-sample variation consistent with previously identified spatiotemporal ecological patterns. This suggests that eDNA sampling from commercial vessels is capable of yielding high quality data relevant for biodiversity surveys and spatial ecological research.

### Advantages of sampling from commercial vessels

Scaling marine eDNA studies to support surveys over large spatial scales, or increasing the temporal frequency of sampling efforts is limited by logistical and financial constraints arising from the high costs and access to survey vessels for use offshore (Bani et al., 2020; Sigsgaard et al., 2020). Using commercial vessels as a platform potentially provides the following advantages: 1) Access to remote offshore locations: commercial vessels make regular regional or transoceanic voyages, travelling 100s or 1000s of kilometres offshore. Open sea areas are often of high biological relevance but are difficult to access with the smaller vessels commonly available to researchers, or which require large dedicated oceanography/fisheries research vessels with high operating costs (Abdulla et al., 2009); 2) Repeatability: commercial vessels, and ferries in particular, follow specific routes with high traffic frequency, thus allowing repeated sequential sampling along the same tracks. In principle this allows for transect based study designs, and facilitates temporal comparisons ranging from days to seasons and years for specific locations; 3) Easy diurnal sampling: commercial vessel scheduling allows for routine operation at night, which may not be feasible for smaller port-based research vessels that have operating restrictions during hours of darkness; 4) Synchronicity: different routes can be surveyed at once, allowing co-ordinated simultaneous sampling over ocean basins; 5) Linear sampling: sampling from vessels allows collection of eDNA over tracks of 3-4 km (versus single point sampling), which may be advantageous for some applications, by increasing the amount and diversity of eDNA recovered; 6) ‘Emission free’ surveys: sampling takes place as an addition to existing journeys, rather than as specifically commissioned research voyages, therefore no extra emissions are attributable to the sampling procedure; 7) Cost effectiveness: commercial vessel platforms can drastically cut the expense of eDNA sampling, since they remove the need to operate dedicated research vessels, and operators may be willing to host researchers and equipment at no or nominal cost; 8) Increased public awareness around conservation issues: participation in eDNA surveys offers opportunities for scientific outreach activities on marine conservation with ferry (vessel) companies and, in turn, the wider public.

### Methodological considerations

For convenient acquisition of eDNA samples from large vessels, we advocate using engine cooling-water taken from the external environment. Most large vessels will have water intakes which can be intercepted by researchers. This can provide samples ‘on demand’, while the vessel is underway, without the need for deployment of additional external equipment. We suggest a Standard Operational Procedure (SOP; see Box 1) employing aluminised plastic storage bags (Bag-in-Box Sampling System (BiBSS); Box 2), for convenient water sampling and storage. Allowing the intake water to flow for 5-8 min before the actual collection ensures that collections will reflect the eDNA profile at the intended point of sampling. Samples are derived from water collected along the track for the duration of the collection window, which may extend for several kilometres at typical vessel cruising speeds. The collection window length is determined by the vessel speed, the flow rate of water drawn from the cooling system, and the speed at which this fills the BiBSS. Adjusting the flow rate may allow for samples to be collected over shorter or longer track lengths, depending on study requirements. In the current study, the water intake was situated 4.5 m below the surface. Vessels with larger drafts will have correspondingly deeper intakes, but the intake position limits sampling to surface waters, and therefore is less flexible than sampling at different depths during a dedicated research cruise. The importance of this constraint will depend on the study research focus, and behaviour of taxa of interest. However, in the current study we detected eDNA from species from multiple depth zones. The extent to which this represents mixing of water across depth strata, and the mechanisms which influence the distribution of eDNA remain to be explored.

#### Box 1. Standard Operational Procedure (SOP) for commercial vessel transect eDNA sampling

- 1) **VOLUME OF SAMPLED WATER**. It would be good practice to filter large volumes of marine waters (up to reach the membrane saturation), in order to retain as much eDNA as possible. Such a volume is however variable, depending on filter characteristics and on water density (e.g. day-time samples saturate the filters quickly, being rich in phytoplankton). According to our experience, maturated in this study and in the analysis carried out in controlled environment (Valsecchi et al. 2020), we suggest the processing of **4-5 litres of marine water per filter**.
- 2) **FILTER POROSITY**. We did not find any significant difference between the three tested NC (nitrocellulose) filter types with porosity 0.22 μm, 0.45 μm, 0.8 μm. However, we suggest to exclude the 0.22 μm pore-size membrane, as filtration is very slow, and saturation is reached after 2-3 litres, without providing a better quantity/quality eDNA. Between the two remaining filter types, we recommend the use of **0.45 μm pore-size membranes**, in order to retain the smallest biological particles, consistently with findings by Li et al. (2018).
- 3) **NUMBER OF REPLICAS** sample replicas are necessary for both a) increase the total amount of eDNA retrieved from each single sampling station (useful for future analyses) and b) reduce the false positive and negative rate inbuilt in the metabarcoding technique (Ficetola et al. 2015). Thus, a **minimum of 3 replicas per station** is advisable (meaning a total of 12-13 litres collected from each sampling station).
- 4) **SAMPLE CONTAINER**. The **Bag in the Box sampling system (BiBSS)** presents many advantages for the collection/preservation/storage of marine water samples for eDNA surveys (see Box S2).
- 5) **SAMPLING STATION DESIGN**. The selection of the geographic positions where establishing FSSs, the fix-sampling stations invariable over cruises, should aim at: 1) **sample spots of biological interest** based on previous observational/literature data; 2) **prioritize points on bathymetric maps indicating habitat changes** (e.g. edge of continental shelf); 3) **select roughly equidistant sampling sites** (about 35-45 nautical miles apart) along the designated shipping lanes, in such a way of both covering the whole route and foreseeing the collection of night samples too. For the same reason, in order to sample adjacent points at different time of the day, it is recommendable to number the stations following the sampling chronological order, meaning that on the map they will not appear in a consecutive order. For example, if 6 fix sampling stations (FSSs) are selected, and 3 will be sampled on the outward journey and 3 on the return journey, the order along the route, on the map, will be: PortA-FSS6-FSS 1-<u>FSS 5</u>-FSS 2-<u>FSS 4</u>-FSS 3-PortB, with the three underscored sampling stations surveyed in the return journey.
- 6) **TIME BEFORE FILTRATION**. Preferably the water contained in the BiB bags should be filtered **immediately after collection**, also to facilitate the sample transportation and avoid the displacement of bulky water samples. However, if this is not possible, a sample storage time of 1-2 weeks between collection and filtering seems to be well tolerated, provided that the BiBs are kept at 4 degrees and away from exposure to the sun during transport. It is important to note that water samples should never be frozen to avoid breaking of cellular components that would result in the release of extracellular DNA which is more easily lost in filtration.
- 7) **TIME BEFORE EXTRACTION**. After filtration **filters should be frozen a.s.a.p**.. The time before extraction does not seem to have a negative effect within the tested time interval, although it is advisable to perform DNA extraction a.s.a.p. after filtration.
- 8) **IMPORTANCE TO COMBINE MOLECULAR TO VISUAL/TAXONOMIC CENSUS**. Until eDNA methodology won’t reach high resolution standards, it is **crucial to combine it with both visual observations (in order to continuously monitor and perfectionate its efficacy) and the taxonomic identification of new species**, followed by the sequencing of their mitogenome in order to fill reference sequences gaps in the molecular databases.

#### Box 2. Advantages of the Bag-in-Box Sampling System (BiBSS) in marine eDNA surveys.

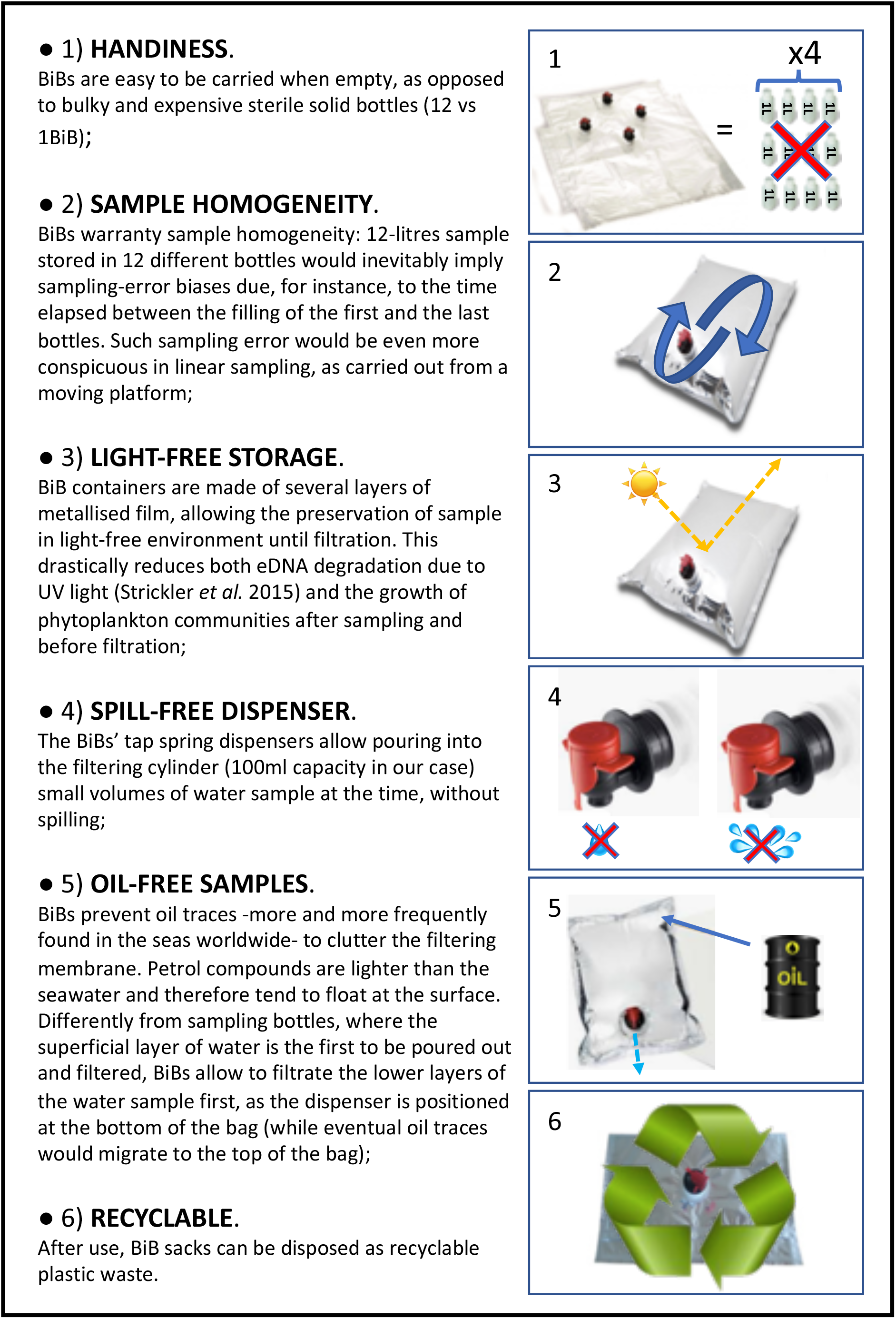

### Marker choice

The taxonomic focus of this study was on marine vertebrates, which, as intermediate and top level predators, are important indicators of the status of marine ecosystems (Hazen et al., 2019). We employed two universal marine vertebrate primer sets, recently developed by our team (Valsecchi et al., 2020). This work reports their first use with environmental samples collected at sea, and corroborates their previous successful validation with marine aquarium samples (Valsecchi et al., 2020). Both MarVer1 (12S rDNA, targeting a similar region to the MiFish primers (Miya et al., 2015)) and MarVer3 (16S rDNA) simultaneously recovered fish and cetacean amplicons, but the taxonomic coverage of the two loci was not fully overlapping. MarVer3 detected almost twice as many teleost species as MarVer1, but not all species found with MarVer1 were also detected by MarVer3 (Table 1, 2; Figure 2, 4). However, there was good concordance between the two markers for the most abundant MOTUs.

Both primers successfully detected cetacean species (Figure 5), but only striped dolphins were consistently detected concurrently. MarVer1 identified sperm whale presence in sample LiGA2.4, while fin whales were identified by both loci, but in non-overlapping samples. This suggests that while both markers have the capability to amplify all four cetacean species, as with other studies, stochasticity still influences taxon detection, reinforcing the importance of replication with a combination of primer sets (Shelton et al., 2016; Sawaya et al., 2019; Djurhuus et al., 2020).

The success of MOTU annotation in all metabarcoding studies is dependent on the accuracy and coverage of reference databases for the chosen loci (Hestetun et al., 2020). In this study, most gaps in taxonomic overlap between the two loci are likely to be due to lack of reference sequences. The previous *in silico* assessment of MarVer1 and MarvVer3 (Valsecchi et al., 2020), indicated that the present coverage of teleost fish for the MarVer1 region is less extensive than for MarVer3, corresponding with the smaller number of MOTUs assigned for MarVer1. However, our annotation pipeline would assign a MOTU to Genus or Family level, if no match within the homology threshold was found at the species level, providing relevant reference sequences were available. Garfish (*Belone belone*) was found in abundance by MarVer3 in the first cruise (e.g. representing more than 50 % of total reads in sample LiGA1.3; Figure 4), but was absent among MarVer1 MOTUs. There are no deposited sequences for the genus *Belone* for the MarVer1 region, but references for three other genera in the same family (Belonidae) are available. However, these were all Pacific or IndoPacific species (*Ablennes hians, Tylosurus acus melanotus, Strongylura anastomella*) and none of them matched with any of the amplicons produced in our study, suggesting that if *Belone belone* amplicons were actually produced using MarVer1, they would have probably remained unannotated due to the lack of reference sequences for the nominal and related taxa. The rapid uptake of eDNA surveys means that reference database coverage can be expected to improve quickly, but some coverage variation dependent on the popularity of different barcode targets will persist (Porter and Hajibabaei, 2018). Taxonomic biases also exist, with many poorly studied groups continuing to be underrepresented in reference databases (Porter and Hajibabaei, 2018; Wangensteen et al., 2018). Improving the latter point will require support from traditional taxonomic approaches (e.g. morphological descriptions) to link sequence information with species descriptions (Cognato et al., 2020).

Polymorphism levels within barcode target regions can also influence the accuracy of annotation (Alberdi et al., 2018). Where variation among species falls below the threshold for MOTU aggregation (98 % homology for this study, equating to 4 to 5 variable sites), automated annotation may not be able to resolve some taxa. In this study, both MarVer1 and MarVer3 had MOTUs assigned to species not considered to be resident in the Mediterranean (e.g. tuna species other than Atlantic and Albacore) due to this issue, which required additional manual curation to resolve.

Elasmobranchs may be underrepresented in our data. Amplicons from rays accounted for 0.04 and 0.033 % of total reads for MarVer1 and MarVer3 respectively, and there were no detections of reads from shark species. These results are consistent with other recent studies which suggest cartilaginous fish species are hard to detect as their eDNA is rare enough to be overwhelmed by teleost eDNA when using universal primers (Stoeckle et al., 2020). The role of eDNA concentration in elasmobranch detection has also been highlighted from aquarium studies where both MarVer1 and MarVer3 detected all the three cartilaginous fish species swimming in the shark tank, while failing to detect the three benthic elasmobranch species (the southern and round stingrays and the longcomb sawfish) (Valsecchi et al., 2020). Elasmobranch detection would probably be facilitated by the use of specific primers for cartilaginous species.

### Spatiotemporal ecological signatures in eDNA profiles

Despite a small sample size, this study detected inter-sample variation consistent with spatiotemporal ecological patterns known from conventional survey approaches. A well-recognised behaviour of bathypelagic species is the diel migrations made as they rise from depth to feed in surface waters at night, particularly in association with lunar cycles (Shima and Swearer, 2019). We detected increased abundance of reads from bathypelagic taxa, notably lantern fish (family Myctophidae) in night samples from Cruise 2, which was close to the full moon (Table S2). This behaviour would also predict increased species diversity in nocturnal samples as bathypelagic species enter the surface layer, mixing with resident taxa. We observed increased alpha diversity for night time samples with both loci, although the difference was not statistically significant (possibly due to low power arising the limited sample size). Night time samples also had significantly higher read abundance for both markers, potentially reflecting increased biomass and eDNA load in water collected at night. This suggests that circadian vertical movements of organisms and/or water masses can bring molecular signals from lower layers to surface waters, and that in some cases sampling of surface waters alone may allow eDNA to characterise community composition representative of different habitats (Suter et al., 2020; West et al., 2020). Thus, it is advisable to collect both day and night-time samples for offshore surveys. This also highlights the capability of eDNA to detect real time changes in biodiversity profiles, reflecting aspects of species behaviour, like diel cycles, which change on short time scales.

The majority of reads for both loci were derived from teleost fish species and overall species composition were consistent with taxa expected for the Mediterranean. Noticeable heterogeneity was observed among sites, and temporal samples at the same location, despite the study covering a relatively small geographic area (approximately 300 km transect length), and close sampling time intervals (3 collections spaced 15 days apart in June-July). Consistent with a previous study in the Bay of Biscay, (Fraija□Fernández et al., 2020), reads from anchovy (*Engraulis* sp.) and sardine (*Sardina* sp.) taxa were the most abundant MOTUs. These species also showed a striking temporal pattern in relative abundance with the former being predominant in the first cruise (18th-19th June 2018) and the latter dominating in the two July cruises (2nd-3rd and 16th-17th July 2018). Mediterranean anchovy populations are known to preferentially spawn with SST below 25°C matching the significant negative correlation observed for *Engraulis* read count abundance and SST in this study (Palomera et al., 2007). Reads from the Mediterranean rainbow wrasse (*Coris julis*), also showed a strong spatiotemporal pattern, being detected at high abundance in night-time samples from station 4 on cruises 2 and 3 (Figure 4). *Coris julis* is typically described as a benthic species so a high abundance in pelagic samples is potentially paradoxical; however, the species’ eggs are pelagic and peak spawning occurs in June coincident with our survey period (Quignard and Pras, 1986; Alonso-Fernández et al., 2014). These observations suggest that some of the strongest fish signals we detected could be related to spawning events rather than to schooling fish (Sigsgaard et al., 2017; Sevellec et al., 2020). It is unclear whether this could also be the case also for the *Belone belone* peak detected in the June cruise, as it is epipelagic and its spawning time is also compatible with the period of our sampling campaign (Tsikliras et al., 2010).

### Cetofauna detection

To date, marine mammal sequences have been detected only occasionally within eDNA samples screened for fish (Closek et al., 2019), or with vertebrate specific primers (Port et al., 2016) or a combination of both (Sigsgaard et al., 2020). Other eDNA studies targeting cetaceans using species-specific primers suggest that cetacean eDNA is hard to detect, even in close spatial or temporal proximity to the source animal. For example, the molecular detection of harbour porpoise (*Phocoena phocoena*) eDNA was observed to diminish at distances greater than 10 meters from a netted sea pen hosting four harbour porpoises, and in the open sea, harbour porpoises were detected in only one of the 24 samples, collected in eight sites known to be attended via a static acoustic survey (Foote et al., 2012). Killer whale (*Orcinus orca*) eDNA was detected in water samples collected from the wake of killer whale pods up to two hours after the encounter (Baker et al., 2018). However, more than half of the sampled encounters produced no or only weak detections, indicating that detection probabilities were low for samples not collected in close temporal association to the animals (Baker et al., 2018). Similarly, bowhead whale (*Balaena mysticetus*) eDNA was found to diminish substantially in water samples collected as little as 10 minutes after the whale presence (Székely et al., 2021). The reported challenges in detecting cetacean eDNA suggest that the results from our pilot study using novel primers optimised to for cetaceans are encouraging (Valsecchi et al., 2020), given that a) they allow for amplification of all vertebrate taxa, meaning low-prevalence eDNA could be overwhelmed by high abundance targets (Rojahn et al., 2021); and b) metabarcoding is less sensitive to detecting rare signals than species specific qPCR assays (Harper et al., 2018; Qu and Stewart, 2019).

Our survey detected four common cetacean species present in the Mediterranean at multiple sites across all three cruises, associated with both visual sighting events and fixed stations. While the limited sample sizes for sampling stations and cruises do not permit formal statistical comparison, trends in the patterns of read abundances were consistent with some reported aspects of cetacean ecology. Firstly, mean read abundances were greater in sighting samples for the two dolphin species, suggesting a potential influence on detection rates of spatiotemporal proximity to animals, fitting those reported in earlier studies (e.g. Székely et al., 2021). Across sites, mean cetacean read abundances were greater for fixed stations 3 and 4 compared to stations 1 and 2. Stations 3 and 4 have bathymetry greater than 200 m and lie close to the continental shelf edge, as did the sighting locations. Across cruises, Cruise 2 had the highest cetacean read counts. Cruise 2 coincided with a near full moon and returned the highest MOTU alpha diversity for both markers. Finally, the highest read abundances for fin and sperm whale were from night-time samples from fixed stations on Cruise 2, and mean abundances for the dolphin species were higher at night when sighting samples were excluded.

Cetacean distributions are closely aligned with prey distributions. Presence in deeper waters close to continental shelves, are a frequently observed pattern, as nutrient rich upwellings can drive concentration of prey species (Shaff and Baird, 2021). Similarly, diurnal variation in activity use of different depth zones, sometimes interacting with the lunar phase, has also been reported (Shaff and Baird, 2021). Increased use of surface waters (<20m) by fin whales (*Balaenoptera physalus*) in the Southern California Bight at night has been observed (Keen et al., 2019). It is noteworthy that lantern fish form a significant component of Mediterranean striped dolphin diet (Dede et al., 2016), with the Madeira lantern fish (*Ceratoscopelus maderensis*) being the second most abundant prey recorded (after *Diaphus* spp, also belonging to Myctophidae family). *Ceratoscopelus maderensis, Hygophum hygomii* and *Hygophum benoiti* lantern fish reads were extremely high on the night of July 2^nd^ 2018, coincident with when the largest numbers of cetacean reads were recorded. These potential links between eDNA and environmental covariates are tantalising and deserve deeper exploration in future work. These results also suggest that in some cases, taking into account target species ecology, such as increased night-time activity, or responses to lunar phase, may help cetacean eDNA researchers enhance detection rates.

## Conclusion and future prospects

This study demonstrates the feasibility of sampling from commercial vessels (Mediterranean ferries) while underway as a strategy to support replicable, systematic marine eDNA surveys in locations that would normally be challenging and expensive for researchers to access. The ISPRA FLT Med cetacean monitoring network currently covers 12 fixed routes around the Mediterranean Sea during all seasons. Therefore, the methodological approach could be extended to other seasons and areas of the Mediterranean basin. eDNA samples are in principle straight forward to collect, meaning that non-specialist professionals, or amateur researchers have potential to contribute to such studies via ‘citizen science’ initiatives, opening the possibility to expand biodiversity surveys across large sea areas and to create the first international marine Mediterranean eDNA bank. This could establish an archive of material for future studies, supporting current work on biodiversity monitoring, and research targeted at specific species of commercial and conservation interest (e.g. Mediterranean monk seal, *Monachus monachus* (Valsecchi E., *preprint*)), and invasive species monitoring. This novel approach for the study of the Mediterranean seascape, will bridge existing biodiversity gaps (e.g. low coverage of pelagic marine communities) and contribute to wildlife management and marine conservation planning. For example, most newly described endemic, cryptogenic or alien species detected in the Mediterranean have been from coastal waters (Chartosia et al., 2018; Katsanevakis et al., 2020). This notion could imply that the number of invasive species in the Mediterranean is currently underestimated and future research needs to focus on offshore waters.

The approach presented in this pilot study could also be extended to any large vessel and to any sea area. While our study focused on Mediterranean ferries, other commercial vessel types such as container ships, tankers, and passenger liners could all offer similar sampling opportunities on any shipping route globally. The high volume and global distribution of commercial vessel traffic mean that such platforms could make an important contribution to the future of ocean monitoring though eDNA, especially with automation of sampling and analysis workflows.

## Supporting information

10 supplementary items

## ACKNOWLEDGEMENTS

We greatly thank Ummey Hany from the Leeds Institute of Molecular Medicine sequencing facility for support on preparation of NGS libraries. We are grateful to Corsica and Sardinia Ferries, in the person of Cristina Pizzutti, for welcoming our proposal of attempting eDNA sampling from their fleet, with a special thanks to the whole crew of the MegaExpress 3 for its availability and support. RL was able to carry out the lab work at Leeds University thanks to the ERASMUS+ programme (2017-1-IT02-KA103-035644). The study is part of the project MeD for Med (Marine eDNA for the Mediterranean), supported also by Bicocca the University of the Crowdfund (grant 2020-CONT-0312).

## LITERATURE CITED

EU Copernicus Marine Service. Global ocean 1/4° physics analysis and forecast updated daily. https://resources.marine.copernicus.eu/?option=com_csw&view=details&product_id=GLOBAL_ANALYSISFORECAST_PHY_CPL_001_015 Accessed: 07/10/2020. .

EU Copernicus Marine Service. Global ocean biogeochemistry analysis and forecast. https://resources.marine.copernicus.eu/?option=com_csw&view=details&product_id=GLOBAL_ANALYSIS_FORECAST_BIO_001_028 Accessed: 05/10/2020.

EU Copernicus Marine Service. Global ocean OSTIA sea surface temperature and sea ice analysis. https://resources.marine.copernicus.eu/?option=com_csw&view=details&product_id=SST_GLO_SST_L4_NRT_OBSERVATIONS_010_001 Accessed 01/10/2020. .

Abdulla, A., Gomei, M., Hyrenbach, D., Notarbartolo-di-Sciara, G., and Agardy, T. (2009). Challenges facing a network of representative marine protected areas in the Mediterranean: prioritizing the protection of underrepresented habitats. ICES Journal of marine science 66(1), 22-28. doi: https://doi.org/10.1093/icesjms/fsn164.

Alberdi, A., Aizpurua, O., Gilbert, M.T.P., and Bohmann, K. (2018). Scrutinizing key steps for reliable metabarcoding of environmental samples. Methods in Ecology and Evolution 9(1), 134-147. doi: https://doi.org/10.1111/2041-210X.12849.

Alonso-Fernández, A., Alós, J., and Palmer, M. (2014). Variability in reproductive traits in the sex-changing fish, Coris julis, in the Mediterranean. Mediterranean Marine Science 15(1), 106-114. doi: http://dx.doi.org/10.12681/mms.455.

Arcangeli, A., Marini, L., and Crosti, R. (2013). Changes in cetacean presence, relative abundance and distribution over 20 years along a trans□regional fixed line transect in the Central Tyrrhenian Sea. Marine Ecology 34(1), 112-121. doi: https://doi.org/10.1111/maec.12006.

Baker, C.S., Steel, D., Nieukirk, S., and Klinck, H. (2018). Environmental DNA (eDNA) from the wake of the whales: droplet digital PCR for detection and species identification. Frontiers in Marine Science 5, 133. doi: https://doi.org/10.3389/fmars.2018.00133.

Bakker, J., Wangensteen, O.S., Chapman, D.D., Boussarie, G., Buddo, D., Guttridge, T.L., et al. (2017). Environmental DNA reveals tropical shark diversity in contrasting levels of anthropogenic impact. Sci Rep 7(1), 16886. doi: https://doi.org/10.1038/s41598-017-17150-2.

Bani, A., De Brauwer, M., Creer, S., Dumbrell, A.J., Limmon, G., Jompa, J., et al. (2020). “Informing marine spatial planning decisions with environmental DNA,” in Advances in Ecological Research. Elsevier), 375–407.

Berry, T.E., Saunders, B.J., Coghlan, M.L., Stat, M., Jarman, S., Richardson, A.J., et al. (2019). Marine environmental DNA biomonitoring reveals seasonal patterns in biodiversity and identifies ecosystem responses to anomalous climatic events. PLoS genetics 15(2), e1007943. doi: https://doi.org/10.1371/journal.pgen.1007943.

Chartosia, N., Anastasiadis, D., Bazairi, H., Crocetta, F., Deidun, A., Despalatovic, M., et al. (2018). New Mediterranean Biodiversity Records (July 2018). Mediterranean Marine Science 19(2), 398. doi: http://dx.doi.org/10.12681/mms.18099.

Closek, C.J., Santora, J.A., Starks, H.A., Schroeder, I.D., Andruszkiewicz, E.A., Sakuma, K.M., et al. (2019). Marine vertebrate biodiversity and distribution within the central California Current using environmental DNA (eDNA) metabarcoding and ecosystem surveys. Frontiers in Marine Science 6, 732. doi: https://doi.org/10.3389/fmars.2019.00732.

Cognato, A.I., Sari, G., Smith, S.M., Beaver, R.A., Li, Y., Hulcr, J., et al. (2020). The essential role of taxonomic expertise in the creation of DNA databases for the identification and delimitation of southeast Asian ambrosia beetle species (Curculionidae: Scolytinae: Xyleborini). Frontiers in Ecology and Evolution 8, 27. doi: https://doi.org/10.3389/fevo.2020.00027.

Dede, A., Salman, A., and Tonay, A.M. (2016). Stomach contents of by-caught striped dolphins (Stenella coeruleoalba) in the eastern Mediterranean Sea. Journal of the Marine Biological Association of the UK 96(4), 869. doi: http://dx.doi.org/10.1017/S0025315415001538.

Djurhuus, A., Closek, C.J., Kelly, R.P., Pitz, K.J., Michisaki, R.P., Starks, H.A., et al. (2020). Environmental DNA reveals seasonal shifts and potential interactions in a marine community. Nature communications 11(1), 1-9. doi: https://www.nature.com/articles/s41467-019-14105-1.

Ficetola, G.F., Pansu, J., Bonin, A., Coissac, E., Giguet-Covex, C., De Barba, M., et al. (2015). Replication levels, false presences and the estimation of the presence/absence from eDNA metabarcoding data. Molecular ecology resources 15(3), 543-556. doi: https://doi.org/10.1111/1755-0998.12338.

Foote, A.D., Thomsen, P.F., Sveegaard, S., Wahlberg, M., Kielgast, J., Kyhn, L.A., et al. (2012). Investigating the potential use of environmental DNA (eDNA) for genetic monitoring of marine mammals. PloS one 7(8), e41781. doi: https://doi.org/10.1371/journal.pone.0041781.

Fraija-Fernández, N., Bouquieaux, M.C., Rey, A., Mendibil, I., Cotano, U., Irigoien, X., et al. (2020). Marine water environmental DNA metabarcoding provides a comprehensive fish diversity assessment and reveals spatial patterns in a large oceanic area. Ecology and Evolution 10(14), 7560-7584. doi: https://doi.org/10.1002/ece3.6482.

Hale, R., Colton, M.A., Peng, P., and Swearer, S.E. (2018). Do spatial scale and life history affect fish–habitat relationships? Journal of Animal Ecology 88(3), 439-449. doi: https://doi.org/10.1111/1365-2656.12924.

Harper, L.R., Lawson Handley, L., Hahn, C., Boonham, N., Rees, H.C., Gough, K.C., et al. (2018). Needle in a haystack? A comparison of eDNA metabarcoding and targeted qPCR for detection of the great crested newt (Triturus cristatus). Ecology and evolution 8(12), 6330-6341. doi: https://doi.org/10.1002/ece3.4013.

Hazen, E.L., Abrahms, B., Brodie, S., Carroll, G., Jacox, M.G., Savoca, M.S., et al. (2019). Marine top predators as climate and ecosystem sentinels. Frontiers in Ecology and the Environment 17(10), 565-574. doi: https://doi.org/10.1002/fee.2125.

Hestetun, J.T., Bye-Ingebrigtsen, E., Nilsson, R.H., Glover, A.G., Johansen, P.-O., and Dahlgren, T.G. (2020). Significant taxon sampling gaps in DNA databases limit the operational use of marine macrofauna metabarcoding. Marine Biodiversity 50(5), 1-9. doi: https://doi.org/10.1007/s12526-020-01093-5.

Hooker, S.K., Cañadas, A., Hyrenbach, K.D., Corrigan, C., Polovina, J.J., and Reeves, R.R. (2011). Making protected area networks effective for marine top predators. Endangered Species Research 13, 203-218. doi: https://doi.org/10.3354/esr00322.

Hunter, M.E., Meigs-Friend, G., Ferrante, J.A., Kamla, A.T., Dorazio, R.M., Diagne, L.K., et al. (2018). Surveys of environmental DNA (eDNA): a new approach to estimate occurrence in Vulnerable manatee populations. Endangered Species Research 35, 101-111. doi: https://doi.org/10.3354/esr00880.

ISPRA (2015). Technical Annex I for the fixed line transect using ferries as platform of observation monitoring protocol.

Katsanevakis, S., Poursanidis, D., Hoffman, R., Rizgalla, J., Rothman, S.B.-S., Levitt-Barmats, Y.a., et al. (2020). Unpublished Mediterranean records of marine alien and cryptogenic species. BioInvasions Records, 2020, vol. 9, núm. 2, p. 165–182.

Keen, E.M., Falcone, E.A., Andrews, R.D., and Schorr, G.S. (2019). Diel Dive Behavior of Fin Whales (Balaenoptera physalus) in the Southern California Bight. Aquatic Mammals 45(2), 233-243. doi: https://doi.org/10.1578/AM.45.2.2019.233.

Li, J., Lawson Handley, L.J., Read, D.S., and Hänfling, B. (2018). The effect of filtration method on the efficiency of environmental DNA capture and quantification via metabarcoding. Molecular Ecology Resources 18(5), 1102-1114. doi: https://doi.org/10.1111/1755-0998.12899.

Matear, L., Robbins, J.R., Hale, M., and Potts, J. (2019). Cetacean biodiversity in the Bay of Biscay: suggestions for environmental protection derived from citizen science data. Marine Policy 109, 103672. doi: https://doi.org/10.1016/j.marpol.2019.103672.

McKnight, D.T., Huerlimann, R., Bower, D.S., Schwarzkopf, L., Alford, R.A., and Zenger, K.R. (2019). microDecon: A highly accurate read□subtraction tool for the post□sequencing removal of contamination in metabarcoding studies. Environmental DNA 1(1), 14-25. doi: https://doi.org/10.1002/edn3.11.

McMurdie, P.J., and Holmes, S. (2013). phyloseq: an R package for reproducible interactive analysis and graphics of microbiome census data. PloS one 8(4), e61217. doi: https://doi.org/10.1371/journal.pone.0061217.

Menegotto, A., and Rangel, T.F. (2018). Mapping knowledge gaps in marine diversity reveals a latitudinal gradient of missing species richness. Nat Commun 9(1), 4713. doi: https://doi.org/10.1038/s41467-018-07217-7.

Miya, M., Sato, Y., Fukunaga, T., Sado, T., Poulsen, J.Y., Sato, K., et al. (2015). MiFish, a set of universal PCR primers for metabarcoding environmental DNA from fishes: detection of more than 230 subtropical marine species. R Soc Open Sci 2(7), 150088. doi: https://doi.org/10.1098/rsos.150088.

Notarbartolo□di□Sciara, G., Agardy, T., Hyrenbach, D., Scovazzi, T., and Van Klaveren, P. (2008). The Pelagos sanctuary for Mediterranean marine mammals. Aquatic Conservation: Marine and Freshwater Ecosystems 18(4), 367-391. doi: https://doi.org/10.1002/aqc.855.

O’Donell, J.L., Kelly, R.P., Shelton, A.O., Samhouri, J.F., Lowell, N.C., and Williams, G.D. (2017). Spatial distribution of environmental DNA in a nearshore marine habitat. PeerJ 5, e3044. doi: https://doi.org/10.7717/peerj.3044.

Palomera, I., Olivar, M.P., Salat, J., Sabatés, A., Coll, M., García, A., et al. (2007). Small pelagic fish in the NW Mediterranean Sea: an ecological review. Progress in Oceanography 74(2-3), 377–396.

Pawlowski, J., Kelly-Quinn, M., Altermatt, F., Apotheloz-Perret-Gentil, L., Beja, P., Boggero, A., et al. (2018). The future of biotic indices in the ecogenomic era: Integrating (e)DNA metabarcoding in biological assessment of aquatic ecosystems. Sci Total Environ 637-638, 1295-1310. doi: https://doi.org/10.1016/j.scitotenv.2018.05.002.

Port, J.A., O’Donnell, J.L., Romero-Maraccini, O.C., Leary, P.R., Litvin, S.Y., Nickols, K.J., et al. (2016). Assessing vertebrate biodiversity in a kelp forest ecosystem using environmental DNA. Molecular Ecology 25(2), 527-541. doi: https://doi.org/10.1111/mec.13481.

Porter, T.M., and Hajibabaei, M. (2018). Over 2.5 million COI sequences in GenBank and growing. PloS one 13(9), e0200177. doi: https://doi.org/10.1371/journal.pone.0200177.

Qu, C., and Stewart, K.A. (2019). Evaluating monitoring options for conservation: comparing traditional and environmental DNA tools for a critically endangered mammal. The Science of Nature 106(3), 1-9. doi: https://doi.org/10.1007/s00114-019-1605-1.

Quignard, J., and Pras, A. (1986). Labridae. Pp: 919–942 in: Whitehead, PJP, M.-L. Bauchot, J.-C. Hureau, J. Nielson & E. Tortonese. Fishes of the North-eastern Atlantic and the Mediterranean.

Rojahn, J., Gleeson, D.M., Furlan, E., Haeusler, T., and Bylemans, J. (2021). Improving the detection of rare native fish species in environmental DNA metabarcoding surveys. Aquatic Conservation: Marine and Freshwater Ecosystems 31(4), 990-997. doi: https://doi.org/10.1002/aqc.3514.

Sawaya, N.A., Djurhuus, A., Closek, C.J., Hepner, M., Olesin, E., Visser, L., et al. (2019). Assessing eukaryotic biodiversity in the Florida Keys National Marine Sanctuary through environmental DNA metabarcoding. Ecology and evolution 9(3), 1029-1040. doi: https://doi.org/10.1002/ece3.4742.

Sevellec, M., Lacoursière□Roussel, A., Bernatchez, L., Normandeau, E., Solomon, E., Arreak, A., et al. (2020). Detecting community change in Arctic marine ecosystems using the temporal dynamics of environmental DNA. Environmental DNA. doi: https://doi.org/10.1002/edn3.155.

Shaff, J.F., and Baird, R.W. (2021). Diel and lunar variation in diving behaviour of rough-toothed dolphins (Steno bredanensis) off Kaua’i, Hawai’i. Marine Mammal Science. doi: https://doi.org/10.1111/mms.12811.

Shelton, A.O., O’Donnell, J.L., Samhouri, J.F., Lowell, N., Williams, G.D., and Kelly, R.P. (2016). A framework for inferring biological communities from environmental DNA. Ecological Applications 26(6), 1645-1659. doi: https://doi.org/10.1890/15-1733.1.

Shima, J.S., and Swearer, S.E. (2019). Moonlight enhances growth in larval fish. Ecology 100(1), e02563. doi: https://doi.org/10.1002/ecy.2563.

Sigsgaard, E.E., Nielsen, I.B., Carl, H., Krag, M.A., Knudsen, S.W., Xing, Y., et al. (2017). Seawater environmental DNA reflects seasonality of a coastal fish community. Marine Biology 164(6), 128. doi: https://doi.org/10.1007/s00227-017-3147-4.

Sigsgaard, E.E., Torquato, F., Frøslev, T.G., Moore, A.B., Sørensen, J.M., Range, P., et al. (2020). Using vertebrate environmental DNA from seawater in biomonitoring of marine habitats. Conservation Biology 34(3), 697-710. doi: https://doi.org/10.1111/cobi.13437.

Stat, M., Huggett, M.J., Bernasconi, R., DiBattista, J.D., Berry, T.E., Newman, S.J., et al. (2017). Ecosystem biomonitoring with eDNA: metabarcoding across the tree of life in a tropical marine environment. Sci Rep 7(1), 12240. doi: https://doi.org/10.1038/s41598-017-12501-5.

Stoeckle, M.Y., Mishu, M.D., and Charlop-Powers, Z. (2020). Improved Environmental DNA Reference Library Detects Overlooked Marine Fishes in New Jersey, United States. Front. Mar. Sci. 7, 226. doi: https://doi.org/10.3389/fmars.2020.00226.

Suter, L., Polanowski, A.M., Clarke, L.J., Kitchener, J.A., and Deagle, B.E. (2020). Capturing open ocean biodiversity: Comparing environmental DNA metabarcoding to the continuous plankton recorder. Molecular Ecology. doi: https://doi.org/10.1111/mec.15587.

Székely, D., Corfixen, N.L., Mørch, L.L., Knudsen, S.W., McCarthy, M.L., Teilmann, J., et al. (2021). Environmental DNA captures the genetic diversity of bowhead whales (Balaena mysticetus) in West Greenland. Environmental DNA. doi: https://doi.org/10.1002/edn3.176.

Tsikliras, A.C., Antonopoulou, E., and Stergiou, K.I. (2010). Spawning period of Mediterranean marine fishes. Reviews in Fish Biology and Fisheries 20(4), 499-538. doi: https://doi.org/10.1007/s11160-010-9158-6.

Valsecchi, E., Bylemans, J., Goodman, S.J., Lombardi, R., Carr, I., Castellano, L., et al. (2020). Novel universal primers for metabarcoding environmental DNA surveys of marine mammals and other marine vertebrates. Environmental DNA 2(4), 460-476. doi: https://doi.org/10.1002/edn3.72.

Valsecchi E., C.E., Pires R., Parmegiani A., Casiraghi M., Galli P., Bruno, A. (preprint). Newly developed ad hoc molecular assays shows how eDNA can witness and anticipate the monk seal recolonization of central Mediterranean. doi: https://doi.org/10.1101/2021.02.13.431078.

Wangensteen, O.S., Palacín, C., Guardiola, M., and Turon, X. (2018). DNA metabarcoding of littoral hard-bottom communities: high diversity and database gaps revealed by two molecular markers. PeerJ 6, e4705. doi: https://doi.org/10.7717/peerj.4705.

West, K.M., Stat, M., Harvey, E.S., Skepper, C.L., DiBattista, J.D., Richards, Z.T., et al. (2020). eDNA metabarcoding survey reveals fine□;scale coral reef community variation across a remote, tropical island ecosystem. Molecular ecology 29(6), 1069-1086. doi: https://doi.org/10.1111/mec.15382.

Wetzel, F.T., Bingham, H.C., Groom, Q., Haase, P., Kõ□alg, U., Kuhlmann, M., et al. (2018). Unlocking biodiversity data: Prioritization and filling the gaps in biodiversity observation data in Europe. Biological conservation 221, 78-85. doi: https://doi.org/10.1016/j.biocon.2017.12.024.

Yamamoto, S., Masuda, R., Sato, Y., Sado, T., Araki, H., Kondoh, M., et al. (2017). Environmental DNA metabarcoding reveals local fish communities in a species-rich coastal sea. Sci Rep 7, 40368. doi: https://doi.org/10.1038/srep40368.

