## Supplementary material for "Ferries and environmental DNA: underway sampling from commercial vessels provides new opportunities for systematic genetic surveys of marine biodiversity": 10 supplementary items

### Supplementary information

| | |
|---|---|
| **Supplementary Table S1** | Table of sampling locations and metadata |
| **Supplementary Figure S1** | Figure showing ferry sampling work flow and apparatus/procedure |
| **Supplementary Figure S2** | Workflow used for dealing with non-Med MOTUs |
| **Supplementary Table S2** | Table detailing resolution of ambiguous MOTU assignments |
| **Supplementary Table S3** | MarVer1 read counts |
| **Supplementary Table S4** | MarVer3 read counts |
| **Supplementary Figure S3** | Figures showing correlations for read counts versus sample types |
| **Supplementary Figure S4** | Barcharts of read count versus site and cruise for each locus |
| **Supplementary Figure S5** | Plots for Anchovy/Sardine environmental analyses |
| **Supplementary Figure S6** | Cluster analysis plots |

**Supplementary Table S1.**

Sample site information including Cruise number; sample codes; distance from closest shore (DfS); Bathymetry at sampling site (m); Sea Surface Temperature (SST; °C); Chlorophyll Concentration (CC; $g/m^3$); Salinity (Practical Salinity Units (PSU), (equivalent to g/kg)); Sample type (fixed or collected after visual observations of cetaceans); Date and time of sampling; Diurnal phase (day or night time sample); Moon phase; Sampling duration (minutes); "Sampled track" columns indicate the number of minutes taken to fill in the 13-litres sampling bag (BiBSS), and the corresponding distance covered by the ferry during the sampling procedure (nautical miles and kilometres), assuming an average cruise speed of 27.5 knots.

| Cruise | Sample code | Sample code brief | Latitude | Longitude | Distance from shore (km) | Bathymetry (m) | Sea Surface Temperature (SST; °C) | Chlorophyll Concentration (mg/m³) | Salinity (PSU; g/kg) | Type | Diurnal | Lunar phase (% visible) | Date | Start | End | Sample duration (minutes) | Sample track length (nm) | Sample track length (km) |
|---|---|---|---|---|---|---|---|---|---|---|---|---|---|---|---|---|---|---|
| 1 | 18-LiGA1.1 | 1.1 | 42.919 | 10.018 | 12 | -71 | 23.50 | 0.1511 | 38.280 | Fixed | Night | 24 waxing | 2018-06-18 | 23:25:00 | 23:35:00 | 10 | 5.27 | 8.49 |
| 1 | 18-LiGA1.2 | 1.2 | 40.985 | 9.684 | 8 | -61 | 22.70 | 0.1686 | 38.254 | Fixed | Night | 24 waxing | 2018-06-19 | 05:45:00 | 05:50:00 | 5 | 2.64 | 4.24 |
| 1 | 18-LiGA1.3 | 1.3 | 41.345 | 9.756 | 38 | -569 | 22.70 | 0.1592 | 38.283 | Fixed | Day | - | 2018-06-19 | 12:14:00 | 12:22:00 | 8 | 4.22 | 6.79 |
| 1 | 18-LiGA1.4 | 1.4 | 42.360 | 9.925 | 30 | -855 | 23.50 | 0.1842 | 38.300 | Fixed | Day | - | 2018-06-19 | 14:27:00 | 14:32:00 | 5 | 2.64 | 4.24 |
| 2 | 18-LiGA2.1 | 2.1 | 42.968 | 10.035 | 19 | -87 | 23.27 | 0.1307 | 38.372 | Fixed | Day | - | 2018-07-02 | 17:12:00 | 17:19:00 | 7 | 3.69 | 5.94 |
| 2 | 18-LiGA2.S1 | 2.S1 | 42.292 | 9.902 | 35 | -875 | 23.97 | 0.1390 | 38.328 | Sighting | Day | - | 2018-07-02 | 18:41:00 | 18:44:00 | 3 | 1.58 | 2.55 |
| 2 | 18-LiGA2.S2 | 2.S2 | 41.856 | 9.832 | 37 | -912 | 24.26 | 0.1361 | 38.408 | Sighting | Day | - | 2018-07-02 | 19:38:00 | 19:43:00 | 5 | 2.64 | 4.24 |
| 2 | 18-LiGA2.2 | 2.2 | 40.985 | 9.684 | 8 | -61 | 23.98 | 0.1506 | 38.316 | Fixed | Night | 88 waning | 2018-07-02 | 21:40:00 | 21:43:00 | 3 | 1.58 | 2.55 |
| 2 | 18-LiGA2.3 | 2.3 | 41.345 | 9.756 | 38 | -569 | 24.27 | 0.1371 | 38.348 | Fixed | Night | 88 waning | 2018-07-03 | 00:27:00 | 00:33:00 | 6 | 3.16 | 5.09 |
| 2 | 18-LiGA2.4 | 2.4 | 42.360 | 9.925 | 30 | -855 | 24.88 | 0.1335 | 38.334 | Fixed | Night | 88 waning | 2018-07-03 | 03:07:00 | 03:11:00 | 4 | 2.11 | 3.40 |
| 3 | 18-LiGA3.1 | 3.1 | 42.931 | 10.026 | 12 | -76 | 25.65 | 0.0952 | 38.416 | Fixed | Day | - | 2018-07-16 | 17:11:00 | 17:13:00 | 2 | 1.05 | 1.70 |
| 3 | 18-LiGA3.S1 | 3.S1 | 42.317 | 9.903 | 35 | -878 | 25.95 | 0.1376 | 38.404 | Sighting | Day | - | 2018-07-16 | 18:38:00 | 18:40:00 | 2 | 1.05 | 1.70 |
| 3 | 18-LiGA3.S2 | 3.S2 | 42.013 | 9.860 | 28 | -769 | 25.38 | 0.1407 | 38.437 | Sighting | Day | - | 2018-07-16 | 19:22:00 | 19:24:00 | 2 | 1.05 | 1.70 |
| 3 | 18-LiGA3.2 | 3.2 | 40.985 | 9.684 | 8 | -61 | 25.23 | 0.1215 | 38.335 | Fixed | Night | 12 waxing | 2018-07-16 | 21:44:00 | 21:46:00 | 2 | 1.05 | 1.70 |
| 3 | 18-LiGA3.3 | 3.3 | 41.345 | 9.756 | 38 | -569 | 23.94 | 0.1334 | 38.312 | Fixed | Night | 12 waxing | 2018-07-17 | 00:33:00 | 00:36:00 | 3 | 1.58 | 2.55 |
| 3 | 18-LiGA3.4 | 3.4 | 42.360 | 9.925 | 30 | -855 | 25.50 | 0.1462 | 38.404 | Fixed | Night | 12 waxing | 2018-07-17 | 03:13:00 | 03:15:00 | 2 | 1.05 | 1.70 |

**Supplementary Figure S1.**

Pictures of some of the project phases: a) Corsica and Sardinia Ferries' Mega Express Three, used as sampling platform; b) visual survey by FLT network member; c) collection of a marine water samples from dedicated pipe in the ferry's engine room; 4) Bag-in-Box Sampling System (BiBSS): the 13L-marine water sample is all contained within the same container.

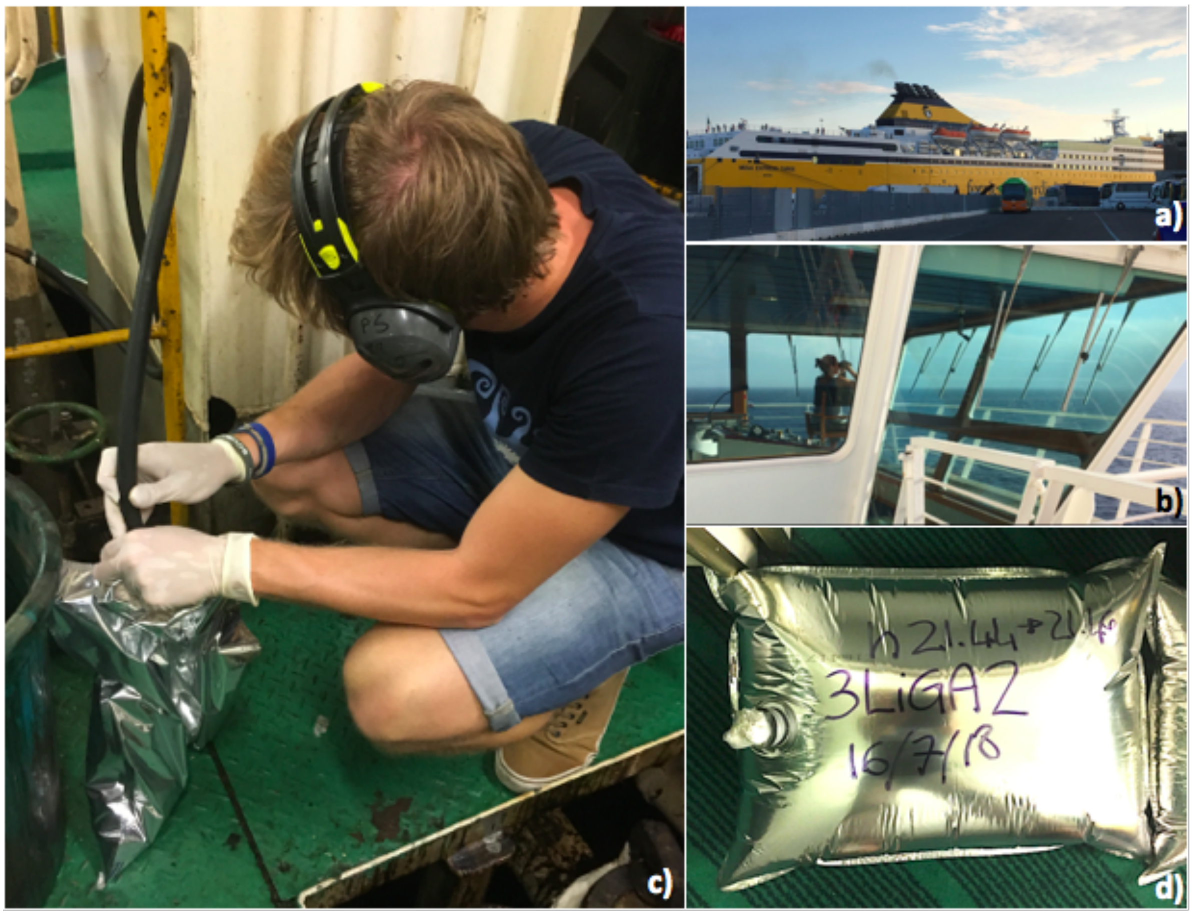

**Supplementary Figure S2** – Workflow used for dealing with non-Med MOTUs. Each detected MOTU not corresponding to known Mediterranean species was subjected to one or both of the questions indicated in the light-blue colour boxes. Depending on the binary response, the availability of reference sequences on GenBank and the degree of differentiation from the most related Mediterranean taxa it is possible to identify seven possible scenarios, four of which (1, 2, 4 and 5) were found in our sample (see Supplementary Table S2).

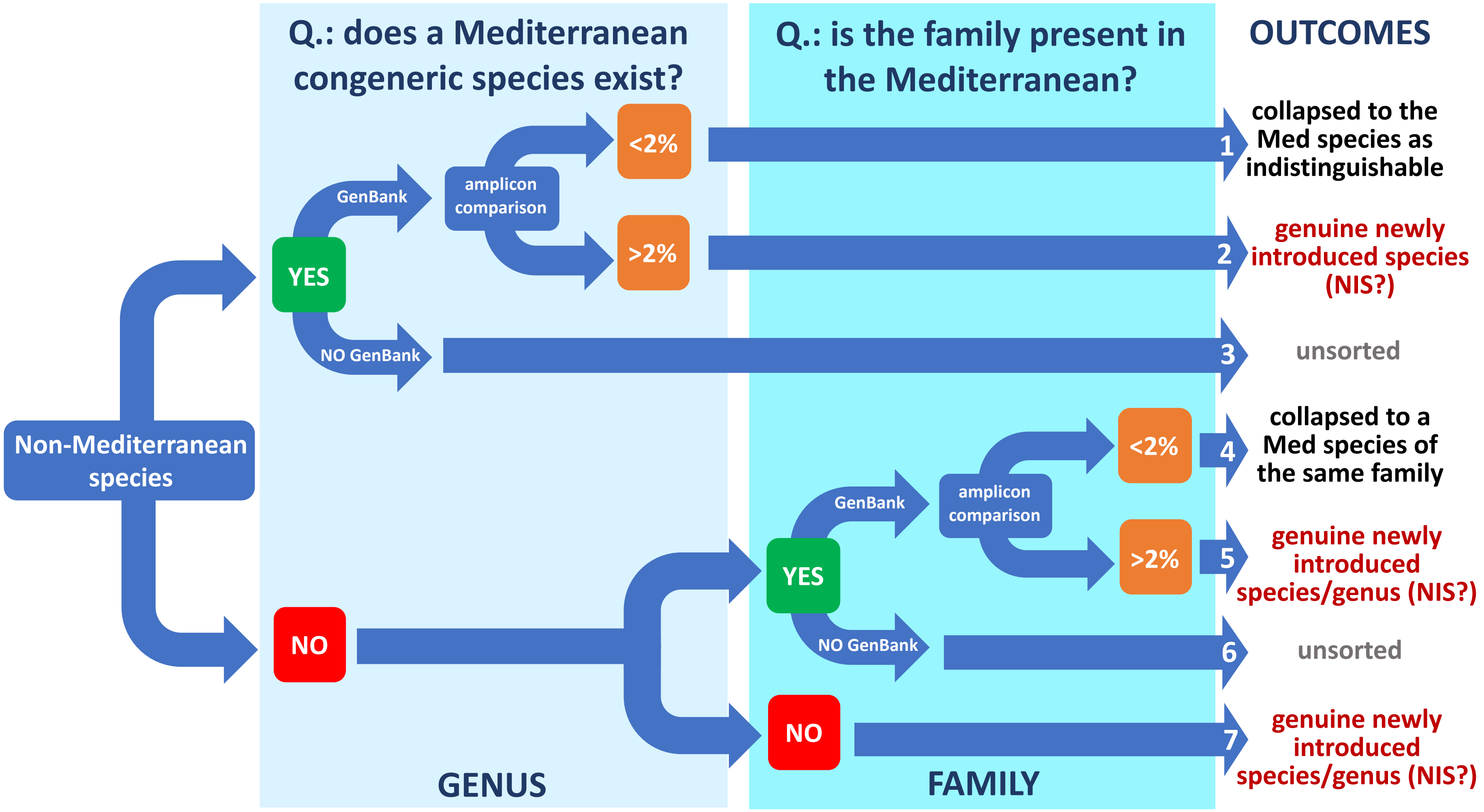

**Supplementary Table S2.** Resolution of annotated MarVer1 and MarVer3 MOTUs assigned to species not recorded as being present in the Mediterranean Sea. Teleost fish MOTUs were compared against the Mediterranean species list from FishBase, and to each comparison was attributed one of the seven scenarios depicted in Figure S2*.

| | **MarVer1: Non-Med MOTUs** | **Congeneric species (or of the same Family) in Mediterranean Sea** | **MOTU Resolution** | ***In silico* molecular comparison annotated and resident MOTUs** | **Scenario (Fig. S2)** |
|---|---|---|---|---|---|
| **1** | *Auxis thazard* | *Auxis rochei* | *Auxis rochei* | *A.rochei:A.thazard* = 2 VS | 1 |
| **2** | *Cheilopogon arcticeps* | *Cheilopogon exsiliens, Cheilopogon heterurus* | *Cheilopogon exsiliens* OR *Cheilopogon heterurus* | *C.arcticeps:C.exsiliens* = 1VS; *C.arcticeps:C. heterurus* = 1VS | 1 |
| **3** | *Chromis alta* | *Chromis chromis, Chromis viridis* | *Chromis chromis* | *C.alta:C.chromis* = 4 VS; ***C.alta:C.viridis* = 10 VS; *C.viridis:C.chromis* = 12 VS** | 1 |
| **4** | *Cololabis saira* | No congeneric: *Scomberesox saurus* (Family – Scomberesocidae) | *Cololabis saira* OR undeposited Scomberesocidae | *Cololabis saira* : *Scomberesox saurus* = **8VS** | 5 |
| **5** | *Dentex tumifrons* | *Dentex dentex, Dentex gibbosus, Dentex macrophthalmus, Dentex maroccanus* | *Dentex tumifrons* | ***D.tumifrons:D.dentex* = 16 VS; *D.tumifrons:D.gibbosus* = 8 VS** | 2 |
| **6** | *Engraulis japonicus* | *Engraulis encrasicolus, Engraulis albidus* | *Engraulis* ssp | ***E.japanicus:E.encrasicolus* = 5VS** | 2 |
| **7** | *Euthynnus affinis* | *Euthynnus alletteratus* | *Euthynnus alletteratus* | *E.affinis:E.alletteratus* = 1VS | 1 |
| **8** | *Gasterochisma melampus* | No congeneric: *Thunnus albacares* (Family – Scombridae) | *Gasterochisma melampus* OR undeposited Scombridae | ***Gasterochisma melampus:Thunnus albacares* = 7 VS** | 5 |
| **9** | *Liza richardsonii* | *Liza aurata* (synonymous *Chelon auratus*) | *Liza aurata* (synonymous *Chelon auratus*) | *L.aurata:L.richardsonii* = 3VS | 1 |
| **10** | *Parargyrops edita* | No congeneric: *Dentex gibbosus, Pagrus pagrus* (Family - Sparidae) | *Dentex gibbosus* | ***Parargyrops.edita:Dentex gibbosus* =5VS** | 4 |
| **11** | *Sardinella longiceps* | *Sardinella madeirensis, Sardinella aurita* | *Sardinella aurita* | ***S.longiceps:S.maderensis* = 26VS;** *S.longiceps:S.aurita* = 1VS*on partial sequence | 1 |
| **12** | *Scomber japonicus* | *Scomber colias, Scomber scombrus* | *Scomber colias* | *S.japanicus:S.colias* = 1VS; ***S.japanicus:S.scombrus* = 9VS** | 1 |
| **13** | *Thunnus albacares* | *Thunnus alalunga, Thunnus thynnus* | *Thunnus* ssp | *T.albacares:T.alalunga* = 2 VS; *T.albacares:T.thynnus* = 2 VS | 1 |
| **14** | *Thunnus maccoyii* | *Thunnus alalunga, Thunnus thynnus* | *Thunnus* ssp | *T.maccoyii:T.alalunga* = 2 VS; *T.maccoyii:T.thynnus* = 2 VS | 1 |
| **15** | *Thunnus orientalis* | *Thunnus alalunga, Thunnus thynnus* | *Thunnus* ssp | *T.orientalis:T.alalunga* = 2 VS; *T.orientalis:T.thynnus* = 4 VS | 1 |
| **16** | *Thunnus tonggol* | *Thunnus alalunga, Thunnus thynnus* | *Thunnus* ssp | *T.tonggol:T.alalunga* = 2 VS; *T.tonggol:T.thynnus* = 3 VS | 1 |
| **17** | *Trachurus japonicus* | *Trachurus trachurus, Trachurus mediterraneus, Trachurus picturatus* | *Trachurus trachurus* | *T.japonicus:T.trachurus* =1 VS | 1 |
| **18** | *Larus glaucoides* | *Larus argentatus* | *Larus argentatus* | *L.glaucoides:L.argentatus*=0 VS | 1 |

| | MarVer3: Non-Med MOTUs | Congeneric species in Mediterranean Sea | MOTU Resolution | *In silico* molecular comparison annotated and resident MOTUs | Scenario |
|---|---|---|---|---|---|
| **1** | *Allothunnus fallai* | No congeneric: *Katsuwonus pelamis* (Family – Scombridae) | *Katsuwonus pelamis* | *Allothunnus fallai:Katsuwonus pelamis* = 4VS | 1 |
| **2** | *Auxis thazard* | *Auxis rochei* | *Auxis rochei* | *A.rochei:A.thazard* =1VS | 1 |
| **3** | *Cyclothone atraria* | *Cyclothone microdon, Cyclothone pygmaea, Cyclothone braueri* | *Cyclothone atraria* | ***C.atraria:C.microdon* = 43 VS, *C.atraria:C.pygmaea* = 39 VS** | 2 |
| **4** | *Dentex canariensis* | *Dentex dentex, Dentex gibbosus, Dentex macrophthalmus, Dentex maroccanus* | *Dentex canariensis* | ***D.canariensis:D.dentex* =12 VS, *D.canariensis:D.gibbosus* =11 VS, *D.canariensis:D.macrophthalmus* =20 VS, *D.canariensis:D.maroccanus* =16 VS, *D.canariensis:D.tumifrons* =25 VS** | 2 |
| **5** | *Engraulis australis* | *Engraulis encrasicolus, Engraulis albidus* | *Engraulis encrasicolus* | *E. australis:E.encrasicolus* = 0 VS | 1 |
| **6** | *Engraulis japonicus* | *Engraulis encrasicolus, Engraulis albidus* | *Engraulis encrasicolus* | *E.japanicus:E.encrasicolus* = 2 VS | 1 |
| **7** | *Lutjanus fulvus* | *Lutjanus argentimaculatus, Lutjanus jocu* | *Lutjanus fulvus* | ***L.fulvus:L.argentimaculatus* = 21 VS; *L.fulvus:L.jocu* = 18 VS** | 2 |
| **8** | *Microstomus kitt* | No congeneric: *Pleuronectes platessa* (Family – Pleuronectidae) | *Pleuronectes platessa* | *Microstomus kitt:Pleuronectes platessa* = 1VS | 1 |
| **9** | *Sardinella longiceps* | *Sardinella aurita, Sardinella maderensis* | *Sardinella aurita* | *S.longiceps:S.aurita* = 0 VS; ***S.longiceps:S.maderensis* = 25 VS** | 1 |
| **10** | *Scomber japonicus* | *Scomber colias, Scomber scombrus* | *Scomber colias* | *S.japanicus:S.colias* = 0 VS*;* ***S.japanicus:S.scombrus* = 8VS** | 1 |
| **11** | *Thunnus albacares* | *Thunnus alalunga, Thunnus thynnus* | *Thunnus ssp* | *T.albacares:T.alalunga* = 3 VS; *T.albacares:T.thynnus* = 0 VS | 1 |
| **12** | *Thunnus atlanticus* | *Thunnus alalunga, Thunnus thynnus* | *Thunnus ssp* | *T.atlanticus:T.alalunga* = 3 VS; *T.atlanticus:T.thynnus* = 2 VS | 1 |
| **13** | *Thunnus maccoyii* | *Thunnus alalunga, Thunnus thynnus* | *Thunnus ssp* | *T.maccoyii:T.alalunga* = 1 VS; *T.maccoyii:T.thynnus* = 1 VS | 1 |
| **14** | *Thunnus obesus* | *Thunnus alalunga, Thunnus thynnus* | *Thunnus ssp* | *T.obesus:T.alalunga* = 2 VS ; *T.obesus:T.thynnus* = 1 VS | 1 |
| **15** | *Trachurus declivis* | *Trachurus trachurus, Trachurus mediterraneus, Trachurus picturatus* | *Trachurus trachurus* | *T.declivis:T.trachurus* = 4 VS; *T.declivis:T.japonicus* = 0 VS | 1 |
| **16** | *Trachurus japonicus* | *Trachurus trachurus, Trachurus mediterraneus, Trachurus picturatus* | *Trachurus trachurus* | *T.japonicus:T.trachurus* = 4 VS; *T.japonicus:T.declivis* = 0 VS | 1 |
| **17** | *Stenella frontalis* | *stenella.coeruleoalba* | *Stenella coeruleoalba* | *S.frontalis:S.coeruleoalba* = 2 VS | 1 |
| **18** | *Tursiops aduncus* | *tursiops.truncatus* | *Tursiops truncatus* | *T.aduncus:T.truncatus* = 1VS | 1 |

*MOTUs were resolved as follow:

1. MOTU were resolved to a specific Mediterranean species if the level of genetic variability (number of variable sites, nVS) was compatible with the 2% threshold of tolerance for MOTU annotation to deposited reference sequences (i.e. nVS<4 and nVS<5 for MarVer1 and MarVer3 respectively).
2. In those cases where either a) the molecular comparison was not possible due to the lack of reference sequence for the Mediterranean congeneric (species in grey font) or b) multiple congeneric species are resident in the Mediterranean and more than two of those differ for <4VS from the MOTU (e.g. Thunnus), the MOTU was resolved at the genus level.
3. Where no other congeneric was resident the MOTU was resolved to family (green fonts).
4. Where the comparison revealed high degree of differentiation incompatible with relaxed annotation (i.e. nVS>4 and nVS>5 for MarVer1 and MarVer3 respectively, highlighted in bold), MOTUs were considered genuine, detecting species not (yet) recorded in the Mediterranean (scenarios 2 and 5, shown in grey). In this case the assigned MOTU was maintained and discussed in the main text.

**Supplementary Table S3. MarVer1 read counts.** MarVer3 read counts for retained taxa after exclusion of potential contamination. binomial – MOTU (species with * indicate resolved MOTUs with original annotation for species not known to be resident in the Mediterranean – see Table S2); L1.1… - samples, Sample:Control ratio – ratio of mean read counts in marine samples, versus 7 negative PCR controls.

| index | binomial | L1.1 | L1.2 | L1.3 | L1.4 | L2.1 | L2.S1 | L2.2 | L2.S2 | L2.3 | L2.4 | L3.1 | L3.S1 | L3.2 | L3.S2 | L3.3 | L3.4 | Total reads | Taxonomy | Sample: Control ratio |
|---|---|---|---|---|---|---|---|---|---|---|---|---|---|---|---|---|---|---|---|---|
| 1 | calonectris.diomedea | 0 | 0 | 0 | 1 | 0 | 0 | 0 | 0 | 0 | 0 | 0 | 0 | 0 | 0 | 12 | 0 | 13 | aves | Inf |
| 2 | larus.argentus *18 | 0 | 0 | 0 | 0 | 0 | 0 | 0 | 0 | 0 | 0 | 1 | 0 | 0 | 0 | 0 | 0 | 1 | aves | Inf |
| 3 | balaenoptera.physalus | 0 | 0 | 0 | 0 | 0 | 0 | 0 | 0 | 0 | 0 | 0 | 0 | 0 | 0 | 0 | 2 | 2 | cetacean | Inf |
| 4 | physeter.. | 0 | 0 | 0 | 0 | 0 | 0 | 0 | 0 | 0 | 52 | 0 | 0 | 0 | 0 | 0 | 0 | 52 | cetacean | Inf |
| 5 | stenella.coeruleoalba | 0 | 0 | 0 | 0 | 0 | 120 | 0 | 116 | 1 | 39 | 1 | 0 | 46 | 0 | 0 | 4 | 327 | cetacean | 23.84375 |
| 6 | tursiops.. | 0 | 1 | 0 | 0 | 0 | 0 | 0 | 0 | 0 | 1 | 0 | 0 | 0 | 0 | 0 | 0 | 2 | cetacean | Inf |
| 7 | mobula.mobular | 83 | 0 | 0 | 63 | 0 | 8 | 1 | 42 | 10 | 1 | 0 | 0 | 0 | 1 | 0 | 5 | 214 | elasmobranch | Inf |
| 8 | pteroplatytrygon.violacea | 0 | 0 | 0 | 0 | 1 | 0 | 23 | 2 | 0 | 0 | 0 | 0 | 0 | 0 | 0 | 0 | 26 | elasmobranch | Inf |
| 9 | aphia.minuta | 0 | 0 | 0 | 0 | 0 | 0 | 0 | 1 | 0 | 0 | 10 | 0 | 0 | 0 | 1 | 2 | 14 | teleost | Inf |
| 10 | atherina.boyeri | 8 | 0 | 0 | 0 | 0 | 0 | 0 | 0 | 0 | 0 | 0 | 0 | 0 | 0 | 0 | 0 | 8 | teleost | Inf |
| 11 | atherina.hepsetus | 0 | 0 | 0 | 0 | 0 | 0 | 0 | 0 | 0 | 0 | 0 | 0 | 0 | 0 | 11 | 0 | 11 | teleost | Inf |
| 12 | auxis.rochei | 378 | 373 | 52 | 181 | 82 | 251 | 759 | 747 | 584 | 55 | 478 | 123 | 3234 | 1457 | 6038 | 1934 | 16726 | teleost | 9.848755 |
| 13 | auxis.rochei *1 | 0 | 1 | 0 | 0 | 0 | 1 | 1 | 0 | 0 | 0 | 7 | 0 | 9 | 0 | 12 | 5 | 36 | teleost | Inf |
| 14 | buenia.affinis | 0 | 0 | 0 | 0 | 1 | 91 | 0 | 12 | 0 | 0 | 0 | 0 | 0 | 0 | 0 | 0 | 104 | teleost | 45.5 |
| 15 | ceratoscopelus.maderensis | 0 | 2 | 0 | 0 | 0 | 0 | 4 | 1 | 0 | 0 | 0 | 1 | 0 | 0 | 1 | 0 | 9 | teleost | Inf |
| 16 | cheilopogon.. *2 | 0 | 0 | 0 | 0 | 0 | 0 | 0 | 0 | 0 | 0 | 0 | 0 | 0 | 0 | 12 | 0 | 12 | teleost | 5.25 |
| 17 | chromis.chromis *3 | 54 | 2 | 0 | 0 | 0 | 21 | 46 | 0 | 0 | 4 | 0 | 0 | 6 | 1 | 0 | 1 | 135 | teleost | Inf |
| 18 | coris.julis | 4323 | 3377 | 99 | 9 | 423 | 402 | 4638 | 101 | 3161 | 10646 | 65 | 290 | 3745 | 32 | 3477 | 49915 | 84703 | teleost | 37.43188 |
| 19 | coryphaena.hippurus | 1 | 83 | 0 | 0 | 0 | 0 | 0 | 0 | 0 | 0 | 0 | 0 | 0 | 1 | 0 | 0 | 85 | teleost | Inf |
| 20 | dentex.gibbosus *10 | 1 | 0 | 0 | 0 | 0 | 0 | 0 | 0 | 0 | 0 | 0 | 0 | 0 | 0 | 0 | 0 | 1 | teleost | Inf |
| 21 | dentex.tumifrons *5 | 0 | 4 | 0 | 0 | 0 | 0 | 0 | 0 | 0 | 0 | 0 | 7 | 7 | 18 | 0 | 0 | 36 | teleost | Inf |
| 22 | dicentrarchus.labrax | 1471 | 1683 | 2046 | 59 | 344 | 161 | 47 | 21 | 1685 | 30 | 3 | 28 | 238 | 134 | 302 | 604 | 8856 | teleost | 5.765625 |
| 23 | engraulis.. *6 | 6 | 0 | 2 | 1 | 5 | 0 | 1 | 2 | 0 | 1 | 5 | 0 | 2 | 5 | 8 | 9 | 47 | teleost | Inf |
| 24 | engraulis.encrasicolus | 32440 | 9804 | 16450 | 6743 | 12944 | 4299 | 2610 | 1994 | 1500 | 15160 | 10537 | 768 | 16333 | 4048 | 41168 | 4668 | 181466 | teleost | 24.18257 |
| 25 | euthynnus.alletteratus | 1280 | 28 | 1 | 3 | 94 | 246 | 454 | 13 | 6 | 2 | 11 | 0 | 1 | 12 | 12 | 3 | 2166 | teleost | 105.2917 |

| | | | | | | | | | | | | | | | | | | | | |
|---|---|---|---|---|---|---|---|---|---|---|---|---|---|---|---|---|---|---|---|---|
| **26** | euthynnus.alletteratus *7 | 1 | 0 | 0 | 0 | 0 | 0 | 0 | 0 | 1 | 0 | 0 | 0 | 2 | 0 | 3 | 1 | 8 | teleost | Inf |
| **27** | gobius.niger | 0 | 0 | 0 | 1 | 0 | 140 | 0 | 0 | 0 | 0 | 3 | 0 | 0 | 12 | 15 | 3 | 174 | teleost | 25.375 |
| **28** | hygophum.hygomii | 5 | 63 | 478 | 3 | 28 | 94 | 32 | 16 | 4075 | 5281 | 1 | 0 | 1 | 10 | 6 | 19 | 10112 | teleost | 98.31111 |
| **29** | katsuwonus.pelamis | 1 | 0 | 0 | 0 | 1 | 0 | 1 | 0 | 0 | 0 | 0 | 0 | 0 | 0 | 0 | 0 | 3 | teleost | Inf |
| **30** | lampanyctus.crocodilus | 0 | 0 | 0 | 0 | 0 | 0 | 22 | 0 | 0 | 0 | 0 | 0 | 0 | 0 | 0 | 0 | 22 | teleost | Inf |
| **31** | liza.aurata | 0 | 0 | 0 | 0 | 0 | 0 | 1 | 0 | 0 | 0 | 0 | 0 | 0 | 0 | 0 | 0 | 1 | teleost | Inf |
| **32** | liza.aurata *9 | 0 | 0 | 1 | 0 | 49 | 590 | 155 | 50 | 31 | 205 | 0 | 1 | 0 | 2 | 22 | 0 | 1106 | teleost | 120.9688 |
| **33** | mola.mola | 0 | 0 | 87 | 1 | 3 | 0 | 0 | 0 | 0 | 0 | 0 | 0 | 0 | 0 | 0 | 2 | 93 | teleost | 40.6875 |
| **34** | myctophum.punctatum | 0 | 0 | 62 | 13 | 0 | 0 | 0 | 0 | 1 | 3 | 0 | 0 | 0 | 0 | 0 | 0 | 79 | teleost | Inf |
| **35** | oedalechilus.labeo | 0 | 0 | 1 | 11 | 0 | 0 | 0 | 0 | 0 | 0 | 0 | 0 | 0 | 0 | 0 | 0 | 12 | teleost | Inf |
| **36** | pagrus.pagrus | 659 | 10 | 14 | 1 | 2 | 2 | 500 | 0 | 8 | 1 | 0 | 2 | 0 | 2 | 128 | 16 | 1345 | teleost | 196.1458 |
| **37** | pomatomus.saltatrix | 1 | 0 | 0 | 1 | 0 | 0 | 0 | 0 | 0 | 0 | 0 | 0 | 0 | 30 | 2 | 34 | teleost | 7.4375 | |
| **38** | sardina.pilchardus | 1904 | 131 | 347 | 334 | 10217 | 2299 | 4514 | 2192 | 17664 | 3331 | 15247 | 1002 | 10125 | 25985 | 76014 | 15721 | 187027 | teleost | 12.01179 |
| **39** | sardinella.aurita *11 | 33 | 21 | 2 | 88 | 33 | 1 | 533 | 3 | 142 | 3 | 20 | 11 | 485 | 25 | 243 | 605 | 2248 | teleost | 30.73438 |
| **40** | scomber.colias *12 | 484 | 135 | 0 | 1 | 6 | 0 | 20 | 2 | 646 | 10 | 2 | 3 | 15 | 7 | 29 | 1 | 1361 | teleost | 595.4375 |
| **41** | scomber.scombrus | 10 | 199 | 1 | 8 | 428 | 19 | 2443 | 34 | 2165 | 0 | 2 | 295 | 53 | 50 | 12 | 236 | 5955 | teleost | 7.48653 |
| **42** | scomberesocidae *4 | 0 | 0 | 0 | 0 | 0 | 0 | 0 | 0 | 0 | 6 | 0 | 0 | 0 | 0 | 0 | 0 | 6 | teleost | Inf |
| **43** | scombridae *8 | 1 | 0 | 0 | 0 | 0 | 0 | 0 | 0 | 0 | 0 | 0 | 0 | 0 | 0 | 0 | 0 | 1 | teleost | Inf |
| **44** | sparus.aurata | 12 | 5 | 1 | 1 | 5 | 1 | 12 | 1 | 303 | 0 | 2 | 2 | 96 | 14 | 135 | 7 | 597 | teleost | 37.3125 |
| **45** | spicara.maena | 0 | 0 | 0 | 127 | 2 | 0 | 0 | 0 | 0 | 0 | 0 | 0 | 0 | 0 | 0 | 0 | 129 | teleost | 28.21875 |
| **46** | thunnus.. *13 | 24 | 24 | 0 | 1 | 15 | 2 | 20 | 32 | 9 | 10 | 5 | 75 | 49 | 166 | 312 | 49 | 793 | teleost | 19.27431 |
| **47** | thunnus.. *14 | 4 | 2 | 0 | 1 | 1 | 1 | 17 | 4 | 6 | 7 | 2 | 0 | 6 | 5 | 41 | 8 | 105 | teleost | 9.1875 |
| **48** | thunnus.. *15 | 0 | 0 | 1 | 0 | 0 | 1 | 0 | 1 | 2 | 1 | 0 | 0 | 2 | 3 | 2 | 4 | 17 | teleost | Inf |
| **49** | thunnus.. *16 | 0 | 0 | 0 | 0 | 0 | 0 | 4 | 2 | 0 | 3 | 0 | 2 | 1 | 1 | 6 | 1 | 20 | teleost | Inf |
| **50** | thunnus.alalunga | 11 | 59 | 28 | 66 | 15 | 809 | 1580 | 969 | 423 | 2245 | 139 | 52 | 1201 | 1182 | 1024 | 2664 | 12467 | teleost | 41.32055 |
| **51** | thunnus.thynnus | 557 | 940 | 286 | 1436 | 802 | 392 | 4483 | 1140 | 2004 | 741 | 156 | 114 | 3734 | 41 | 5194 | 368 | 22388 | teleost | 47.09014 |
| **52** | trachurus.trachurus *17 | 1058 | 1833 | 712 | 244 | 63 | 220 | 1022 | 35 | 253 | 2575 | 183 | 25 | 1566 | 79 | 1572 | 1122 | 12562 | teleost | 43.967 |
| **53** | xiphias.gladius | 1 | 1 | 32 | 1 | 0 | 556 | 0 | 3 | 1 | 123 | 1 | 0 | 0 | 0 | 0 | 2 | 721 | teleost | 63.0875 |

**Supplementary Table S4.** MarVer3 read counts for retained taxa after exclusion of potential contamination. Binomial – MOTU (species with * indicate resolved MOTUs with original annotation for species not known to be resident in the Mediterranean – see Table S2); L1.1… - samples, Sample:Control ratio – ratio of mean read counts in marine samples, versus 7 negative PCR controls.

| index | Binomial | L1.1 | L1.2 | L1.3 | L1.4 | L2.1 | L2.S1 | L2.2 | L2.S2 | L2.3 | L2.4 | L3.1 | L3.S1 | L3.2 | L3.S2 | L3.3 | L3.4 | Total reads | Taxonomy | Sample: Control ratio |
|---|---|---|---|---|---|---|---|---|---|---|---|---|---|---|---|---|---|---|---|---|
| 1 | balaenoptera.physalus | 0 | 0 | 0 | 0 | 0 | 0 | 90 | 0 | 211 | 0 | 0 | 0 | 0 | 1 | 1 | 0 | 303 | cetacean | Inf |
| 2 | stenella.coeruleoalba | 1 | 2 | 57 | 4 | 5 | 1094 | 120 | 556 | 265 | 421 | 3 | 2 | 7 | 22 | 0 | 6 | 2565 | cetacean | 102.0170 |
| 3 | stenella.coeruleoalba *17 | 0 | 0 | 1 | 0 | 0 | 4 | 0 | 0 | 0 | 2 | 0 | 0 | 0 | 0 | 0 | 0 | 7 | cetacean | Inf |
| 4 | tursiops.truncatus *18 | 0 | 0 | 293 | 6 | 3 | 830 | 4 | 8 | 69 | 14 | 2 | 0 | 3 | 3 | 1 | 0 | 1236 | cetacean | 135.1875 |
| 5 | mobula.mobular | 1 | 0 | 0 | 0 | 2 | 188 | 65 | 82 | 0 | 1 | 1 | 0 | 0 | 0 | 0 | 0 | 340 | elasmobranch | Inf |
| 6 | aglaura.hemistoma | 0 | 0 | 0 | 0 | 0 | 0 | 0 | 1 | 0 | 13 | 0 | 0 | 0 | 0 | 0 | 0 | 14 | invertebrate | Inf |
| 7 | geryonia.proboscidalis | 0 | 0 | 0 | 0 | 1 | 0 | 0 | 0 | 3 | 0 | 0 | 0 | 0 | 0 | 0 | 0 | 4 | invertebrate | Inf |
| 8 | liriope.tetraphylla | 0 | 1 | 0 | 1 | 16 | 20 | 2 | 2 | 13 | 3 | 0 | 0 | 0 | 0 | 3 | 0 | 61 | invertebrate | 26.6875 |
| 9 | phascolosoma.. | 9 | 116 | 0 | 0 | 0 | 0 | 195 | 0 | 173 | 0 | 1 | 2 | 64 | 0 | 0 | 0 | 560 | invertebrate | 245.0000 |
| 10 | apogon.imberbis | 0 | 0 | 0 | 0 | 0 | 0 | 0 | 0 | 0 | 0 | 0 | 0 | 80 | 0 | 0 | 0 | 80 | teleost | Inf |
| 11 | auxis.rochei *2 | 143 | 698 | 57 | 76 | 223 | 139 | 644 | 5797 | 556 | 65 | 224 | 278 | 3537 | 941 | 2595 | 776 | 16749 | teleost | 101.7734 |
| 12 | belone.belone | 1197 | 42470 | 54722 | 4992 | 1558 | 796 | 129 | 127 | 475 | 92 | 78 | 452 | 282 | 525 | 78 | 34 | 108007 | teleost | 55.9870 |
| 13 | belone.svetovidovi | 74 | 7 | 152 | 3 | 9 | 1 | 2 | 3 | 1 | 5 | 1 | 1 | 3 | 9 | 1 | 1 | 273 | teleost | 5.9719 |
| 14 | boops.. | 0 | 0 | 0 | 167 | 1 | 0 | 1 | 0 | 0 | 0 | 0 | 0 | 0 | 0 | 11 | 0 | 180 | teleost | 39.3750 |
| 15 | brama.brama | 0 | 0 | 0 | 0 | 1 | 0 | 3 | 0 | 165 | 0 | 0 | 0 | 0 | 1 | 0 | 0 | 170 | teleost | 74.3750 |
| 16 | buenia.affinis | 1 | 0 | 0 | 0 | 0 | 0 | 54 | 0 | 48 | 0 | 0 | 0 | 0 | 0 | 0 | 0 | 103 | teleost | Inf |
| 17 | ceratoscopelus.maderensis | 37 | 60 | 18 | 12 | 569 | 18 | 18899 | 341 | 7627 | 99 | 68 | 11 | 3323 | 34 | 302 | 170 | 31588 | teleost | 191.9410 |
| 18 | chelon.. | 0 | 0 | 0 | 0 | 75 | 1 | 192 | 13 | 192 | 0 | 0 | 0 | 51 | 1 | 4 | 2 | 531 | teleost | Inf |
| 19 | chelon.labrosus | 0 | 0 | 0 | 0 | 0 | 0 | 0 | 0 | 0 | 0 | 0 | 0 | 0 | 0 | 3 | 0 | 3 | teleost | Inf |
| 20 | chromis.chromis | 5 | 2 | 4 | 0 | 2 | 323 | 743 | 82 | 411 | 10 | 5 | 2 | 137 | 4 | 20 | 1 | 1751 | teleost | 153.2125 |
| 21 | coris.julis | 3846 | 11485 | 130 | 9 | 208 | 304 | 5173 | 377 | 3598 | 14354 | 74 | 235 | 4762 | 51 | 1417 | 29836 | 75859 | teleost | 54.1408 |
| 22 | cyclothone.atraria | 0 | 0 | 0 | 0 | 0 | 0 | 0 | 17 | 0 | 0 | 0 | 0 | 7 | 0 | 4 | 3 | 31 | teleost | Inf |
| 23 | dentex.canariensis | 0 | 0 | 0 | 0 | 2 | 0 | 0 | 1 | 0 | 0 | 0 | 0 | 0 | 3 | 0 | 0 | 6 | teleost | Inf |
| 24 | dentex.maroccanus | 0 | 11 | 0 | 0 | 6 | 0 | 0 | 4 | 0 | 0 | 1 | 13 | 9 | 65 | 0 | 0 | 109 | teleost | 47.6875 |
| 25 | dicentrarchus.. | 389 | 323 | 6 | 1 | 83 | 54 | 8 | 1 | 158 | 2 | 2 | 1 | 7 | 19 | 6 | 96 | 1156 | teleost | 29.7500 |
| 26 | dicentrarchus.labrax | 146 | 37 | 779 | 20 | 47 | 6 | 61 | 35 | 2196 | 7 | 13 | 3 | 123 | 40 | 86 | 12 | 3611 | teleost | 21.0642 |
| 27 | diplodus.annularis | 13 | 2 | 1 | 3 | 13 | 7 | 2212 | 2 | 1166 | 96 | 6 | 0 | 30 | 8 | 15 | 2 | 3576 | teleost | 82.3421 |

| | | | | | | | | | | | | | | | | | | | |
|---|---|---|---|---|---|---|---|---|---|---|---|---|---|---|---|---|---|---|---|
| **28** | engraulis.. | 1755 | 1133 | 792 | 34 | 273 | 349 | 27 | 200 | 21 | 12655 | 52 | 62 | 871 | 40 | 322 | 31 | 18617 | teleost | 50.2774 |
| **29** | engraulis.encrasicolus | 71870 | 37474 | 30372 | 3961 | 24880 | 9778 | 4849 | 7425 | 5186 | 24232 | 2920 | 1715 | 30360 | 2019 | 33724 | 3052 | 293817 | teleost | 49.7465 |
| **30** | engraulis.encrasicolus *5 | 325 | 277 | 255 | 44 | 281 | 40 | 24 | 48 | 24 | 158 | 43 | 39 | 691 | 9 | 225 | 531 | 3014 | teleost | 10.5490 |
| **31** | engraulis.encrasicolus *6 | 3 | 1 | 1 | 2 | 3 | 1 | 0 | 1 | 0 | 1 | 0 | 0 | 9 | 0 | 7 | 4 | 33 | teleost | 14.4375 |
| **32** | euthynnus.alletteratus | 324 | 6 | 0 | 0 | 0 | 2 | 2 | 378 | 0 | 74 | 5 | 0 | 73 | 4 | 14 | 0 | 882 | teleost | 128.6250 |
| **33** | gobius.. | 0 | 0 | 0 | 0 | 0 | 0 | 0 | 0 | 0 | 0 | 0 | 7 | 35 | 0 | 11 | 0 | 53 | teleost | Inf |
| **34** | gobius.fallax | 0 | 0 | 0 | 0 | 0 | 0 | 0 | 0 | 0 | 0 | 0 | 0 | 0 | 0 | 4 | 0 | 4 | teleost | Inf |
| **35** | gobius.niger | 0 | 0 | 3 | 4 | 207 | 0 | 0 | 0 | 0 | 0 | 0 | 0 | 0 | 0 | 0 | 2 | 216 | teleost | 47.2500 |
| **36** | hygophum.benoiti | 2 | 25 | 3 | 8 | 52 | 23 | 175 | 3 | 16782 | 225 | 0 | 0 | 14 | 10 | 0 | 21 | 17343 | teleost | 135.4922 |
| **37** | hygophum.hygomii | 5 | 44 | 89 | 4 | 18 | 100 | 83 | 44 | 6635 | 4512 | 2 | 1 | 8 | 6 | 2 | 13 | 11566 | teleost | 97.3101 |
| **38** | katsuwonus.pelamis *1 | 0 | 0 | 0 | 0 | 0 | 0 | 0 | 1 | 0 | 0 | 0 | 0 | 0 | 0 | 0 | 0 | 1 | teleost | Inf |
| **39** | lampanyctus.crocodilus | 1 | 0 | 0 | 0 | 1 | 3 | 487 | 0 | 2 | 0 | 1 | 0 | 0 | 0 | 0 | 0 | 495 | teleost | Inf |
| **40** | lithognathus.mormyrus | 0 | 2 | 1 | 1 | 0 | 1 | 125 | 2 | 3 | 0 | 7 | 6 | 560 | 0 | 328 | 18 | 1054 | teleost | 92.2250 |
| **41** | lutjanus.fulvus | 1 | 0 | 0 | 0 | 0 | 0 | 0 | 0 | 0 | 0 | 0 | 0 | 0 | 0 | 0 | 0 | 1 | teleost | Inf |
| **42** | millerigobius.macrocephalus | 0 | 0 | 0 | 0 | 0 | 0 | 1 | 0 | 0 | 0 | 0 | 2 | 0 | 0 | 44 | 97 | 144 | teleost | Inf |
| **43** | mola.mola | 0 | 1 | 0 | 1 | 0 | 387 | 0 | 7 | 0 | 1 | 0 | 0 | 0 | 1 | 0 | 0 | 398 | teleost | 43.5313 |
| **44** | mullus.barbatus | 572 | 5 | 77 | 125 | 1860 | 428 | 1275 | 290 | 397 | 173 | 11 | 19 | 89 | 1 | 1604 | 144 | 7070 | teleost | 99.7782 |
| **45** | myctophum.punctatum | 4 | 162 | 1510 | 503 | 44 | 13 | 78 | 1 | 5262 | 270 | 1 | 2 | 8 | 4 | 43 | 13 | 7918 | teleost | 115.4708 |
| **46** | oblada.melanura | 57 | 3864 | 557 | 15 | 18 | 9 | 143 | 9 | 297 | 35 | 1 | 1 | 10 | 10 | 34 | 3 | 5063 | teleost | 52.7396 |
| **47** | odondebuenia.balearica | 1 | 0 | 0 | 0 | 1 | 0 | 1 | 1 | 0 | 1 | 0 | 2 | 0 | 0 | 142 | 3 | 152 | teleost | 66.5000 |
| **48** | pagellus.erythrinus | 618 | 4 | 0 | 2 | 76 | 2 | 182 | 0 | 2 | 0 | 1 | 1 | 0 | 2 | 71 | 15 | 976 | teleost | 85.4000 |
| **49** | platichthys.. | 0 | 0 | 1 | 0 | 0 | 0 | 0 | 0 | 0 | 0 | 0 | 0 | 0 | 0 | 0 | 1 | 2 | teleost | Inf |
| **50** | pleuronectes.platessa *8 | 2 | 1 | 0 | 0 | 1 | 0 | 1 | 0 | 0 | 1 | 0 | 0 | 0 | 0 | 2 | 0 | 8 | teleost | Inf |
| **51** | sardina.. | 4594 | 507 | 543 | 732 | 27998 | 8787 | 9356 | 17237 | 30271 | 4067 | 6426 | 13908 | 14505 | 34647 | 51070 | 10116 | 234764 | teleost | 46.9206 |
| **52** | sardina.pilchardus | 13 | 2 | 2 | 2 | 127 | 38 | 38 | 230 | 122 | 26 | 20 | 17 | 94 | 70 | 161 | 18 | 980 | teleost | 61.2500 |
| **53** | sardinella.aurita | 172 | 629 | 22 | 29 | 684 | 1299 | 15194 | 433 | 8990 | 88 | 222 | 189 | 7286 | 115 | 3178 | 11174 | 49704 | teleost | 79.6538 |
| **54** | sardinella.aurita *9 | 0 | 0 | 0 | 0 | 0 | 0 | 0 | 0 | 1 | 0 | 0 | 0 | 0 | 0 | 0 | 0 | 1 | teleost | Inf |
| **55** | scomber.. | 2 | 126 | 3 | 3 | 170 | 3 | 481 | 88 | 324 | 0 | 0 | 98 | 84 | 19 | 1 | 5 | 1407 | teleost | 123.1125 |
| **56** | scomber.colias *10 | 100 | 71 | 0 | 0 | 41 | 1 | 16 | 1 | 855 | 7 | 0 | 0 | 2 | 0 | 3 | 1 | 1098 | teleost | 96.0750 |
| **57** | scomber.scombrus | 0 | 0 | 0 | 0 | 2 | 0 | 9 | 0 | 3 | 0 | 0 | 0 | 0 | 0 | 0 | 0 | 14 | teleost | Inf |
| **58** | scomberesox.saurus | 0 | 0 | 0 | 0 | 0 | 0 | 0 | 1 | 0 | 79 | 0 | 0 | 0 | 0 | 0 | 0 | 80 | teleost | Inf |

| | | | | | | | | | | | | | | | | | | | | |
|---|---|---|---|---|---|---|---|---|---|---|---|---|---|---|---|---|---|---|---|---|
| **59** | scorpaena.notata | 0 | 0 | 2 | 1 | 141 | 0 | 241 | 13 | 2 | 0 | 899 | 4 | 6 | 0 | 1 | 1 | 1311 | teleost | 573.5625 |
| **60** | scorpaena.porcus | 0 | 0 | 0 | 0 | 0 | 0 | 0 | 0 | 0 | 0 | 0 | 0 | 0 | 0 | 0 | 1 | 1 | teleost | Inf |
| **61** | serranus.atricauda | 0 | 0 | 0 | 0 | 0 | 0 | 1 | 0 | 0 | 0 | 0 | 0 | 0 | 0 | 3 | 0 | 4 | teleost | Inf |
| **62** | serranus.cabrilla | 8 | 0 | 0 | 0 | 0 | 152 | 1 | 2 | 0 | 2 | 1 | 0 | 0 | 0 | 96 | 1 | 263 | teleost | 115.0625 |
| **63** | serranus.hepatus | 0 | 0 | 0 | 0 | 1 | 0 | 73 | 0 | 1 | 0 | 0 | 0 | 0 | 0 | 120 | 4 | 199 | teleost | 43.5312 |
| **64** | serranus.scriba | 6 | 26 | 68 | 2 | 14 | 2 | 3607 | 59 | 541 | 2 | 6 | 6 | 164 | 3 | 341 | 8 | 4855 | teleost | 236.0069 |
| **65** | solea.senegalensis | 2 | 1 | 1 | 1 | 0 | 0 | 4 | 0 | 0 | 0 | 0 | 0 | 0 | 5 | 4 | 0 | 18 | teleost | Inf |
| **66** | sparus.aurata | 0 | 2 | 0 | 0 | 2 | 0 | 2 | 1 | 461 | 0 | 0 | 0 | 15 | 0 | 43 | 1 | 527 | teleost | 115.2812 |
| **67** | spicara.smaris | 0 | 0 | 0 | 0 | 1 | 0 | 159 | 0 | 280 | 0 | 0 | 0 | 0 | 1 | 33 | 0 | 474 | teleost | Inf |
| **68** | symphodus.tinca | 0 | 0 | 0 | 0 | 0 | 0 | 0 | 0 | 0 | 0 | 0 | 0 | 0 | 0 | 2 | 0 | 2 | teleost | Inf |
| **69** | tetrapturus.belone | 0 | 0 | 0 | 0 | 0 | 0 | 0 | 23 | 0 | 0 | 0 | 0 | 0 | 0 | 0 | 0 | 23 | teleost | Inf |
| **70** | thalassoma.pavo | 1 | 0 | 1 | 1 | 3 | 0 | 1044 | 0 | 3 | 3 | 1 | 0 | 20 | 2 | 4 | 5 | 1088 | teleost | Inf |
| **71** | thunnus.. *11 | 241 | 900 | 220 | 333 | 778 | 1793 | 3925 | 5120 | 2876 | 509 | 132 | 63 | 6125 | 129 | 2172 | 102 | 25418 | teleost | 117.0566 |
| **72** | thunnus.. *12 | 0 | 0 | 0 | 0 | 0 | 1 | 2 | 5 | 1 | 0 | 0 | 0 | 3 | 0 | 2 | 0 | 14 | teleost | Inf |
| **73** | thunnus.. *13 | 2 | 9 | 1 | 1 | 4 | 66 | 61 | 1050 | 129 | 68 | 35 | 11 | 55 | 7 | 82 | 78 | 1659 | teleost | 51.8438 |
| **74** | thunnus.. *14 | 8 | 11 | 2 | 7 | 8 | 26 | 76 | 220 | 12 | 3 | 71 | 7 | 74 | 8 | 159 | 15 | 707 | teleost | 30.9312 |
| **75** | thunnus.alalunga | 21 | 89 | 106 | 9 | 16 | 1402 | 640 | 8078 | 337 | 2282 | 104 | 108 | 968 | 461 | 458 | 1046 | 16125 | teleost | 135.6671 |
| **76** | thunnus.thynnus | 0 | 1 | 0 | 0 | 0 | 0 | 4 | 2 | 2 | 2 | 0 | 0 | 15 | 0 | 2 | 0 | 28 | teleost | Inf |
| **77** | trachinus.draco | 0 | 0 | 1 | 2 | 277 | 1 | 0 | 0 | 84 | 0 | 0 | 0 | 1 | 0 | 40 | 2 | 408 | teleost | 59.5000 |
| **78** | trachurus.. | 0 | 4 | 2 | 0 | 0 | 0 | 1 | 0 | 3 | 4 | 0 | 0 | 0 | 0 | 0 | 0 | 14 | teleost | Inf |
| **79** | trachurus.picturatus | 0 | 4 | 1 | 0 | 2 | 0 | 92 | 0 | 5 | 13 | 1 | 0 | 2 | 0 | 2 | 1 | 123 | teleost | Inf |
| **80** | trachurus.trachurus *15 | 0 | 0 | 0 | 0 | 0 | 0 | 0 | 0 | 0 | 0 | 0 | 0 | 0 | 0 | 1 | 0 | 1 | teleost | Inf |
| **81** | trachurus.trachurus *16 | 793 | 4255 | 716 | 107 | 298 | 396 | 2945 | 55 | 1491 | 4267 | 63 | 22 | 2502 | 20 | 643 | 378 | 18951 | teleost | 92.1229 |
| **82** | trigloporus.lastoviza | 0 | 0 | 0 | 0 | 0 | 0 | 0 | 0 | 0 | 0 | 0 | 0 | 0 | 0 | 2 | 0 | 2 | teleost | Inf |
| **83** | xiphias.gladius | 0 | 1 | 1 | 5 | 1 | 989 | 2 | 192 | 2 | 147 | 2 | 0 | 2 | 9 | 0 | 0 | 1353 | teleost | 197.3125 |
| **84** | xyrichtys.novacula | 17 | 2 | 4 | 2 | 11 | 3 | 4933 | 9 | 38 | 3 | 10 | 0 | 1 | 3 | 7 | 0 | 5043 | teleost | 245.1458 |
| **85** | zebrus.zebrus | 0 | 0 | 0 | 0 | 0 | 0 | 0 | 1 | 1 | 0 | 0 | 0 | 47 | 3 | 15 | 3 | 70 | teleost | Inf |
| **86** | zeus.. | 0 | 0 | 0 | 0 | 0 | 0 | 0 | 0 | 0 | 0 | 0 | 0 | 0 | 0 | 1 | 0 | 1 | teleost | Inf |

**Supplementary Figure S3.** Read counts distribution for the two loci MarVer1 and MarVer3 in relation to different parameters: a) time from sample collection to sample filtration (Tf); b) time from end of filtration to DNA extraction (Te); c) membrane porosity; d) day vs nocturnal samples.

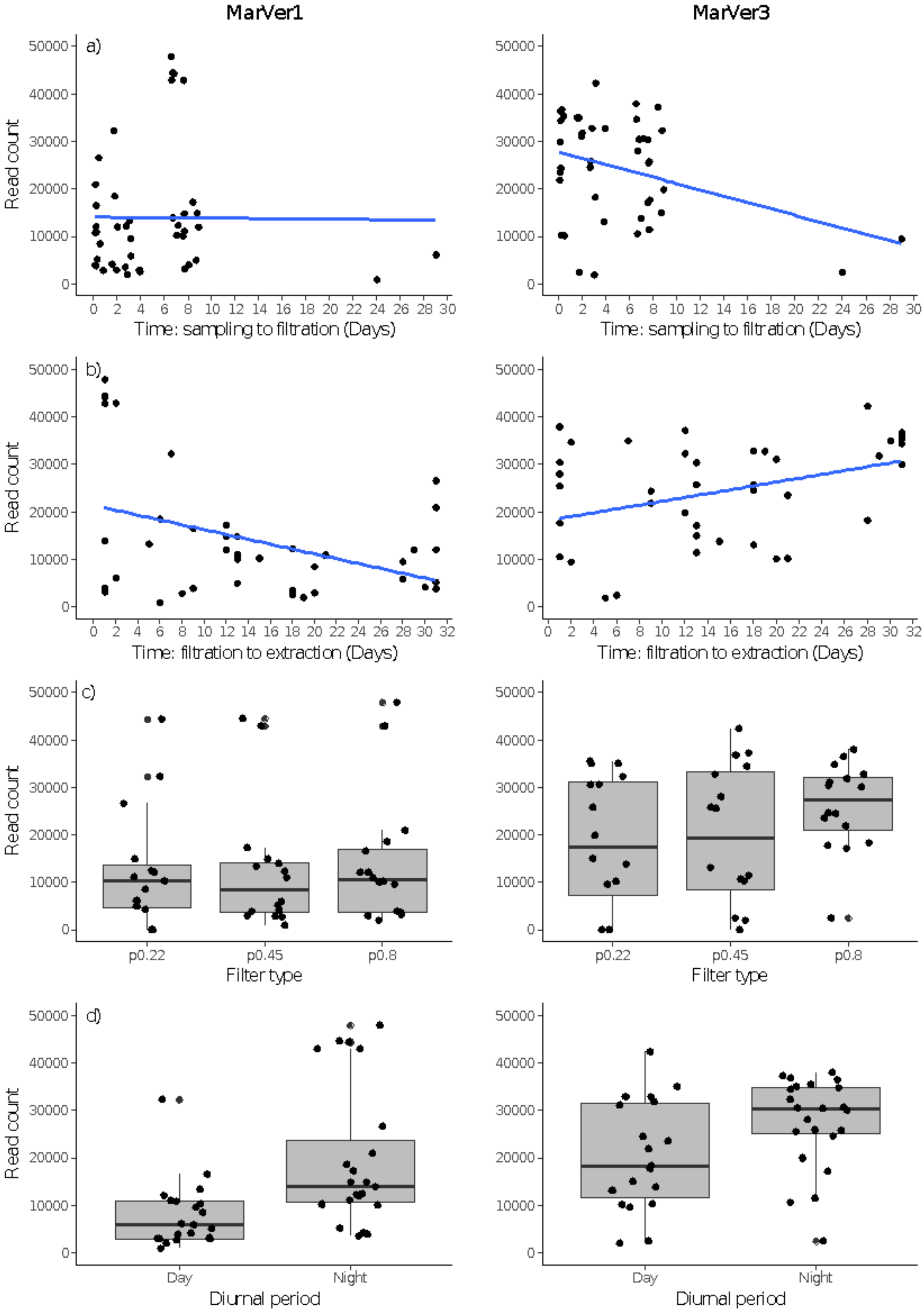

**Supplementary Figure S4**. Barcharts of read count versus site and cruise for each locus.

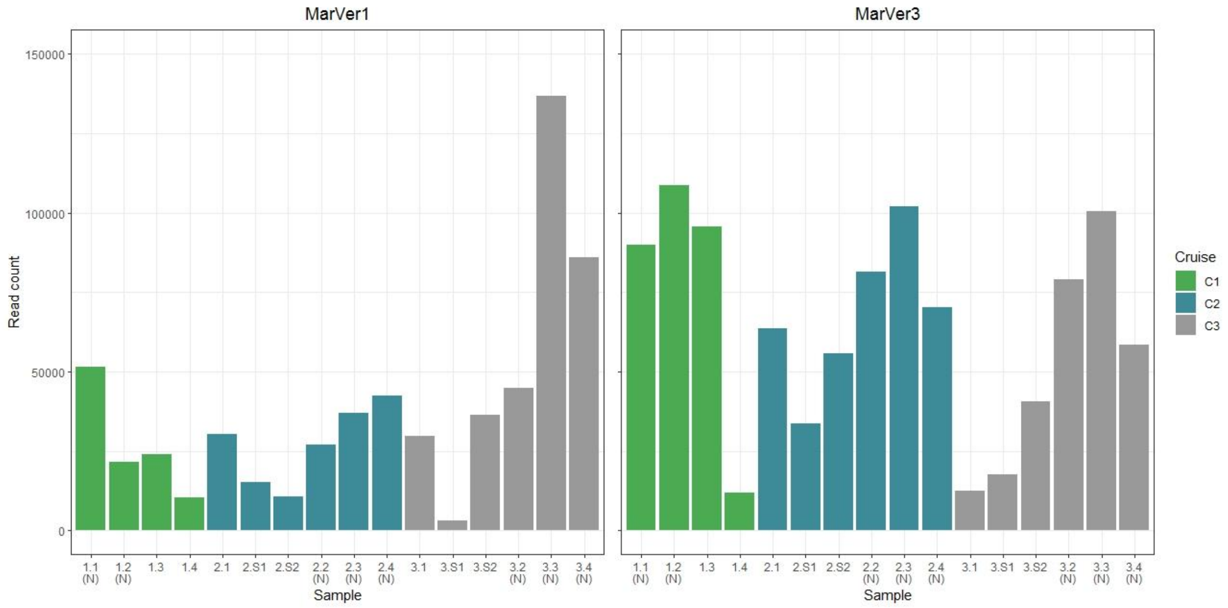

**Supplementary Figure S5**. Plots showing correlation between sample read abundance and sea surface temperature (SST) for anchovy (*Engraulis*), Sardinella, and sardine (*Sardina*) MOTUs, abundance relative to SST and salinity.

### MarVer1

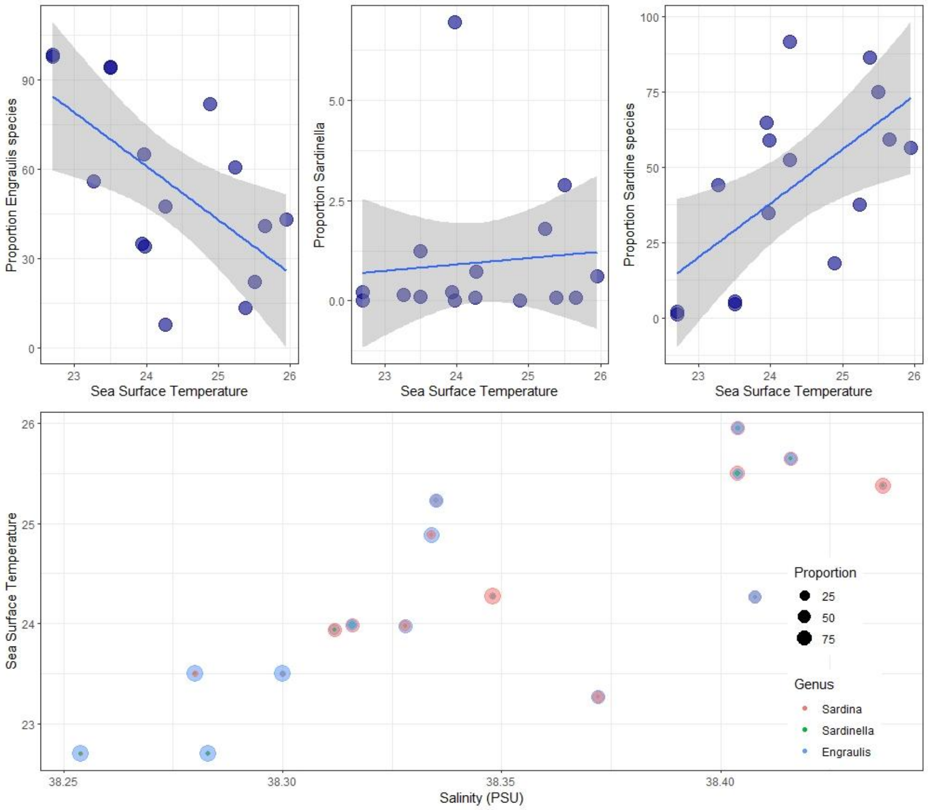


### MarVer3

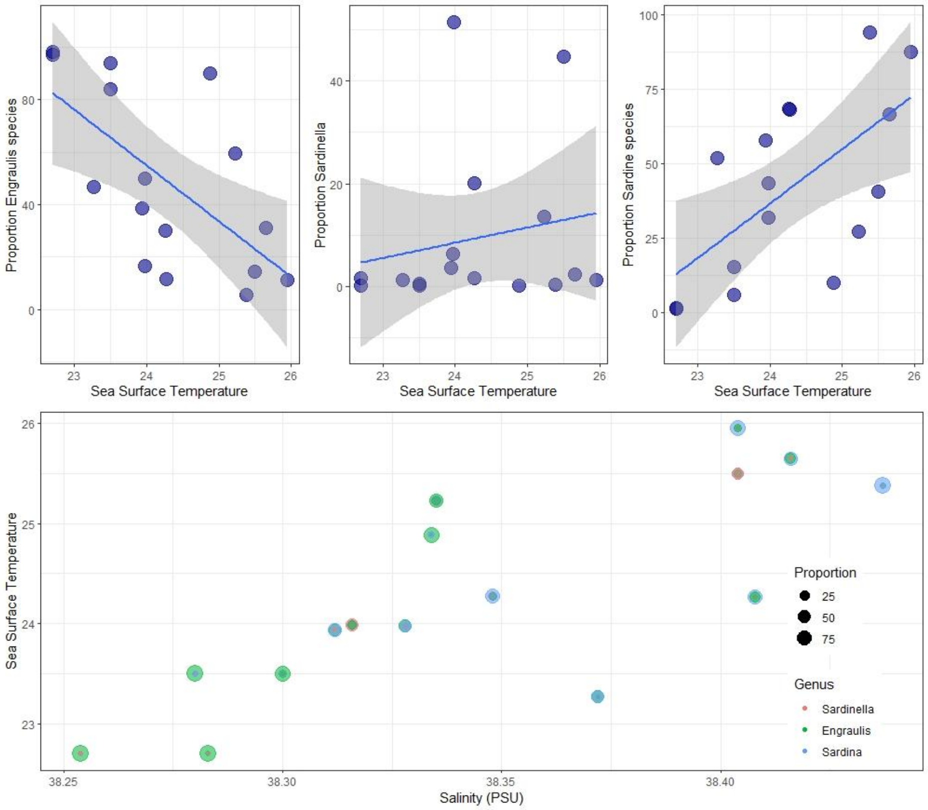

**Supplementary Figure S6**. Hierarchical cluster analysis plots based on Bray-Curtis distance measure of sample similarity for MarVer1 and MarVer3 datasets. (N) – night time sample, (S) cetacean sighting sample.

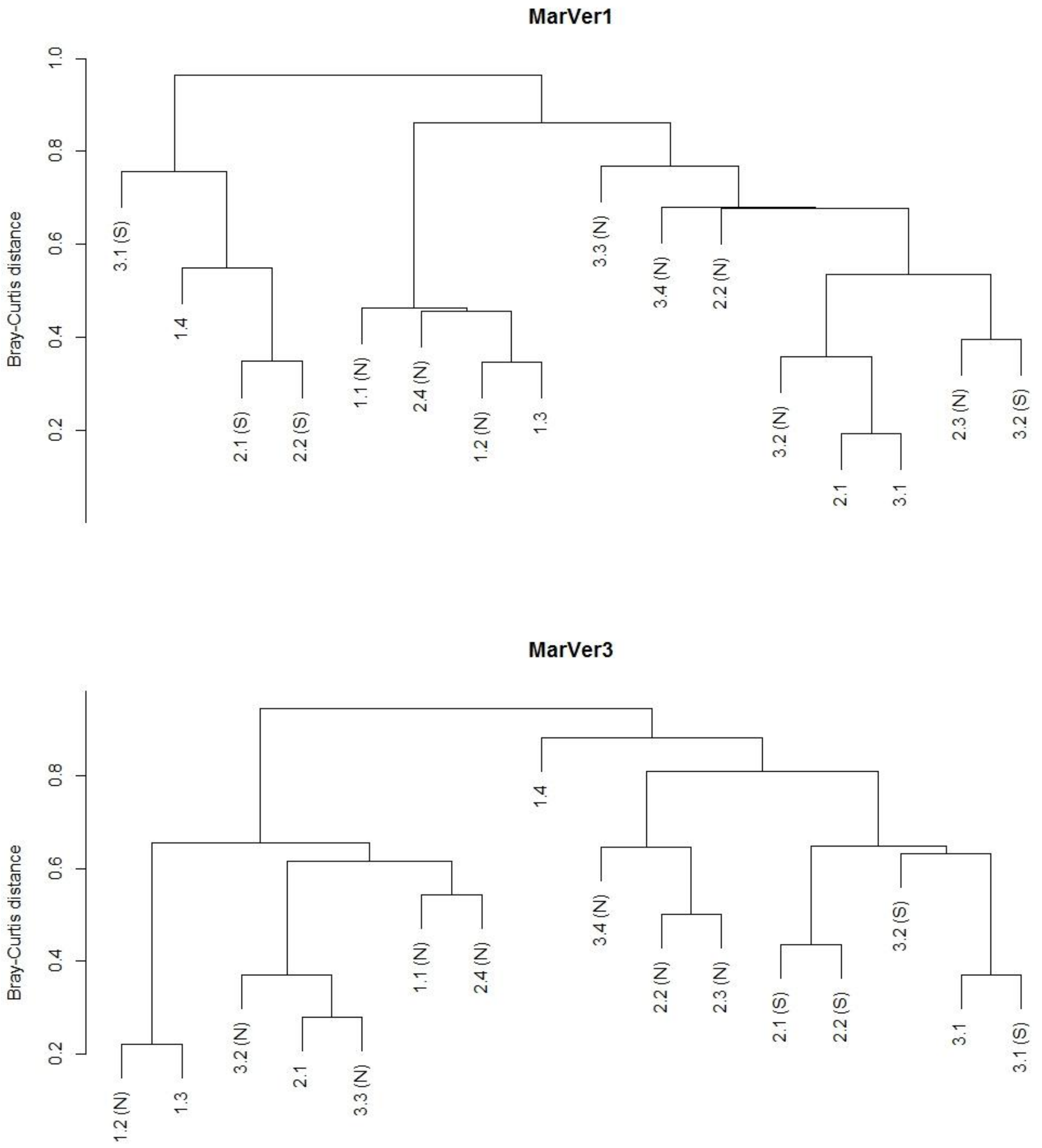
